# A microbial growth-coupled platform for *in vivo* interrogation of Rubisco oxygenase activity

**DOI:** 10.64898/2026.07.10.737725

**Authors:** Enrico Orsi, Kai Kabuth, Simone Cusimano, Mathias Herløv-Wagner, Rutger Verbakel, Francesco Luppino, Michele Partipilo, Benoit de Pins, Elad Noor, Lena Maria Hümmler, Michael Mülleder, Steffen N. Lindner, Markus Ralser, Pablo I. Nikel

**Affiliations:** BRIGHT, Technical University of Denmark, 2800 Kgs Lyngby, Denmark; Department of Biology, University of Naples “Federico II”, Naples, Italy; Department of Plant and Environmental Sciences, Weizmann Institute of Science, Rehovot, Israel; Department of Biochemistry, Charité Universitätsmedizin Berlin, Freie Universität Berlin and Humboldt-Universität, 10117, Berlin, Germany

## Abstract

Rubisco catalyzes the primary CO_2_-fixing reaction of the biosphere, yet its competing oxygenation reaction reduces net global carbon fixation and has resisted direct exploration in living cells. Here, we engineer an auxotrophic *Escherichia coli* strain in which 2-phosphoglycolate, the direct product of Rubisco oxygenation, becomes essential for growth, making bacterial fitness a quantitative proxy for oxygenation flux *in vivo*. This provides direct access to catalytic selectivity, something previously inaccessible to carboxylation-coupled assays. The platform enables screening of phylogenetically diverse Form II Rubisco and phosphoribulokinase (Prk) variants circumventing protein purification and extensive *in vitro* characterization. Adaptive laboratory evolution under oxygenation-selective pressure identified two mutations: Rubisco M115I genetically rebalances the *in vivo* carboxylation/oxygenation trade-off (resulting in 6-fold reduction in *k_cat,_*_C_), while Prk N216T improves overall flux without altering selectivity. This platform makes Rubisco’s least-studied catalytic function selectable and evolvable *in vivo*, opening the carboxylation/oxygenation trade-off to systematic genetic dissection and engineering.

## Introduction

Enzyme promiscuity is widely present in nature and challenges the textbook view of enzymes as exceptionally specific catalysts (1, 2). The most abundant enzyme in the biosphere, ribulose-1,5-bisphosphate (RuBP) carboxylase/oxygenase (Rubisco), with a total mass of approximately 0.7 Gt and responsible for more than 99% of the global CO_2_ fixation (3, 4), is no exception. Alongside its essential carboxylation reaction, Rubisco catalyzes a competing oxygenation that directly limits the efficiency of carbon fixation on a global scale.

Rubisco emerged in a CO_2_-rich atmosphere prior to the Great Oxidation Event. With the rise of oxygenic photosynthesis, it was increasingly challenging to discriminate between CO_2_ and O_2_ as atmospheric CO_2_ declined and O_2_ levels rose (5). Shaped by diverse selective pressures across environmental niches (6), Rubisco has undergone gradual albeit extremely slow improvements in carboxylation efficiency (7). These include for example, enhancements in catalytic kinetics in plants (7); structural modifications such as the recruitment of an additional subunit in Form I Rubisco to improve CO_2_/O_2_ discrimination (8); and the adoption of carbon-concentrating mechanisms in diverse organisms to boost local CO_2_ availability (6, 9).

Despite these evolutionary adaptations, Rubisco retains its oxygenase activity, with oxygenation events exceeding 20% of total catalytic turnovers (5, 10, 11), that significantly reduce net CO_2_ fixation. Oxygenation adds O_2_ to RuBP, producing one molecule of 3-phosphoglycerate (3PG) and one molecule of 2-phosphoglycolate (2PG). The latter is highly reactive and recycled via photorespiration, a process that consumes ATP and reducing power while releasing previously fixed CO_2_, thereby reducing the overall efficiency of carbon fixation (10, 12, 13).

To mitigate the impact of Rubisco oxygenation, synthetic biology efforts have targeted either the enzyme itself through, for example, directed evolution (14–17), or the photorespiration process by introducing bypass routes in metabolism that prevent CO_2_ loss (13, 18–21). Yet, oxygenation remains virtually unavoidable. Rubisco’s kinetic parameters appear constrained by trade-offs among turnover, substrate affinity, and specificity (4, 22–26), and while its oxygenation mechanism is understood (23, 27, 28), studying it has historically relied on laborious *in vitro* approaches that strip the enzyme of its cellular context.

Recently, growth-coupled *in vivo* microbial platforms have begun addressing this limitation for carboxylation, enabling sequence-fitness mapping at scale (17). Growth-coupling offers a route to study enzymes in their physiological context: by tailoring microbial fitness to depend on a specific enzymatic reaction, growth rate becomes a living proxy for catalytic activity *in vivo* (29–31). The same principle extends to continuous selective pressure, enabling directed laboratory evolution under defined metabolic constraints (29, 30, 32–36). For Rubisco specifically, the established Δ*rpiAB* (Δ*rpi*) platform exploits this logic by coupling ribulose 5-phosphate (Ru5P) detoxification to Prk–Rubisco activity (17, 37), but cannot resolve the carboxylation–oxygenation partition that underlies Rubisco’s central catalytic trade-off, because both reactions consume RuBP. Therefore, oxygenase activity has remained an unresolved intrinsic feature rather than a tractable signal. Its physiological consequences and evolutionary potential *in vivo* have therefore not been systematically explored.

Here we developed a growth-coupled platform coupling bacterial fitness directly to Rubisco oxygenation. We repurposed an *E. coli* auxotrophic metabolic sensor originally designed for glycolate sensing (38) to harbor heterologous *prk* and Rubisco genes. In this context, Rubisco oxygenation synthesizes 2PG and, through downstream reactions, replenishes the glycolate pool in the auxotroph. This provides direct access to catalytic selectivity (previously impossible with traditional Rubisco-coupled assays) enabling systematic dissection of sequence determinants controlling oxygenase activity across Rubisco phylogenies. Adaptive laboratory evolution (ALE) identified two mutations with complementary functions: one (located in Rubisco) rebalanced the *in vivo* carboxylation/oxygenation selectivity, while the other (in Prk) enhanced flux irrespective of the selectivity. Altogether, this work establishes a tractable *in vivo* framework for dissecting and modifying Rubisco’s least-studied catalytic function.

## Results

### *In silico* demonstration of growth complementation through Rubisco oxygenation

Building on a compact *E. coli* metabolic model (39), we previously designed mutant strains in which glyoxylate serves as a key metabolic intermediate for engineering glycolate auxotrophies (38). Here, we employ the same metabolic reconstruction to assess if Rubisco oxygenation can complement growth *in silico* in the absence of externally added glycolate.

We selected the auxotrophy design with the highest sensitivity to glycolate i.e., the lowest glycolate flux required to restore growth, corresponding to the combined deletions of isocitrate lyase (Δ*ICL*) and serine hydroxymethyltransferase (Δ*GHMT2r*) (**Fig. 1A**). To this background, which underlies the previously described strains C1+GLY-AUX and GLY-AUX (38), we added BiGG reactions for Prk and for the two Rubisco half-reactions, carboxylation (RBC,_C_) and oxygenation (RBC,_O_). To probe the contribution of each reaction independently of total enzyme activity, we constrained the model so that total Rubisco flux remained constant while the ratio between the two activities was scanned across nine orders of magnitude as 10^φ^ = v(RBC,_O_) / v(RBC,_C_), with φ ∈ [–4, +4].

**Fig. 1.**
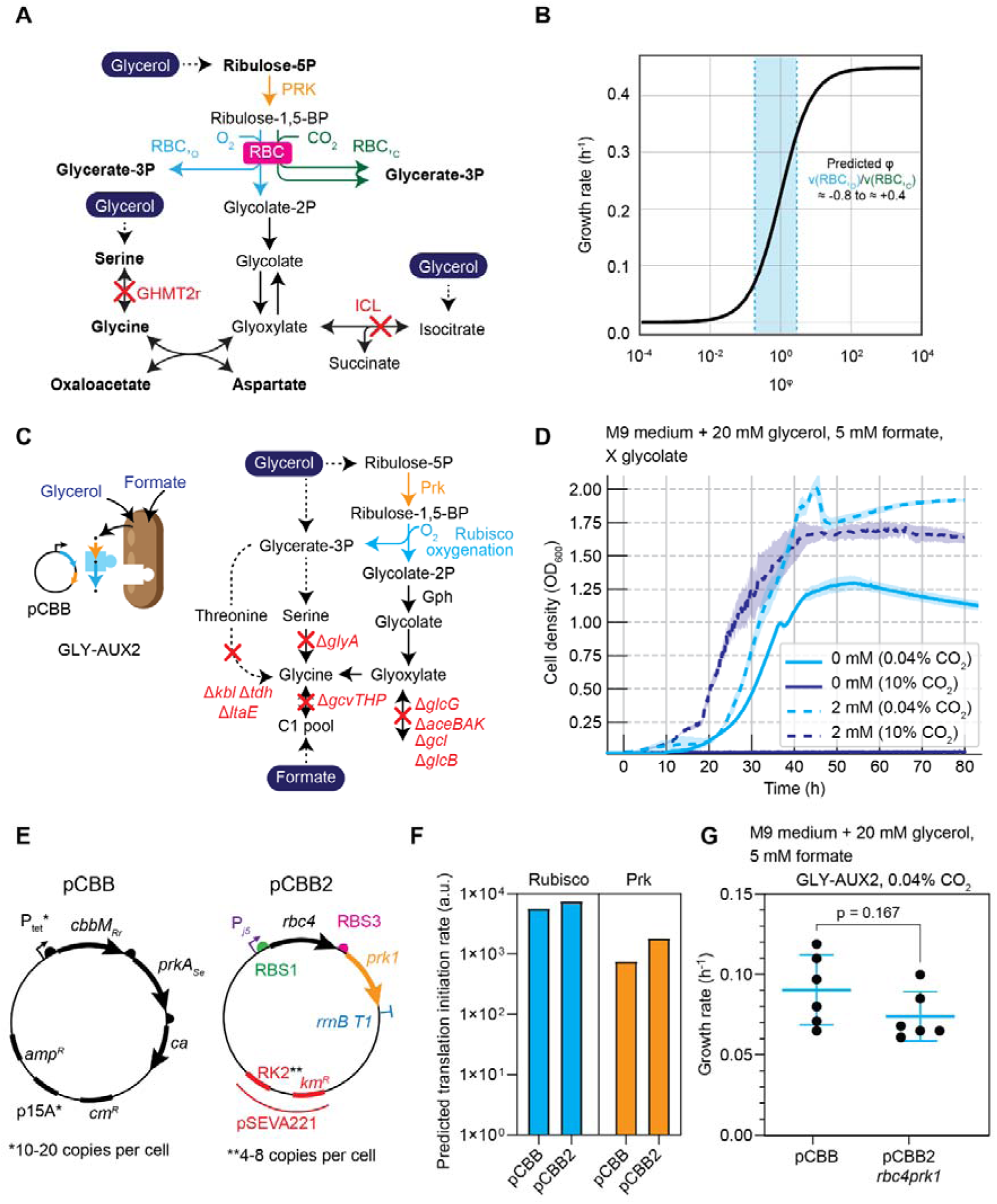
*In silico* and *in vivo* establishment of a growth-coupled sensor for Rubisco oxygenase activity and standardization of the Prk–Rubisco expression system. A) Metabolic network reconstruction of the compact *E. coli* model used in this study, showing the key enzyme deletions (*ICL* and *GHMT2r*) and the heterologous reactions introduced: phosphoribulokinase (PRK) and the two Rubisco half-reactions, carboxylation (RBC_,C_) and oxygenation (RBC_,O_). B) Predicted growth rate as a function of the oxygenation-to-carboxylation flux ratio (φ) in the *in silico* GLY-AUX2 strain. Growth rate increases monotonically with the relative contribution of oxygenation, confirming that the auxotrophic design selectively responds to Rubisco oxygenase flux. C) Schematic representation of GLY-AUX2 and the growth complementation strategy through Prk and Rubisco oxygenase activity. D) Growth profiles of GLY-AUX2 transformed with pCBB under relaxed and selective conditions at ambient CO_2_ (0.04%) and 10% CO_2_. Curves represent one biological replicate measured in technical triplicate. Shaded areas represent standard error with 95% confidence interval. Experiments were performed in biological triplicate. E) Schematic representation of the pCBB architecture and the modifications introduced in pCBB2, shown here carrying *rbc4* and *prk1*. F) Predicted translation initiation rates (TIR) for the Prk and Rubisco expression units in pCBB and pCBB2*rbc4prk1*. G) Growth rates of GLY-AUX2 strains harboring pCBB or pCBB2*rbc4prk1* at ambient CO_2_ (0.04%). Data are mean ± s.d. (n=3 biological replicates); individual points represent technical duplicates. Two-tailed unpaired t-test; p-values indicated.

Under this constraint, the predicted growth rate of the GLY-AUX design rose monotonically with φ, tracking the increasing oxygenation share (**Fig. 1B**). Because the strain is auxotrophic for 2PG-derived metabolites and not for 3PG, its growth is intrinsically insensitive to carboxylation whereas oxygenation sets the predicted growth rate. The model thus supports stoichiometrically that, in this genetic background, growth depends specifically on Rubisco’s oxygenation flux (**Extended Data Fig. 1A**). Using literature values of Rubisco’s *S*_C/O_ and the expected intracellular CO_2_/O_2_ ratio under ambient conditions (**Supplementary File 1**), we estimated a physiologically realistic φ range of −0.8 to +0.4, corresponding to roughly ≈10 to ≈20% of total Rubisco flux going through oxygenation, substantiating that physiologically relevant oxygenation rates are sufficient to complement growth.

To test whether this selectivity is a property of the strain rather than of the Rubisco reactions themselves, we examined the predicted growth response of the established Δ*rpiAB* (Δ*RPI*) sensor background as a function of independently varied carboxylation and oxygenation fluxes (**Supplementary Fig. S1A, C**). In Δ*RPI*, which uses Rubisco as a detoxification route for Ru5P (17, 37), predicted growth was positive across the entire phase plane wherever either flux was nonzero: carboxylation and oxygenation contributed equivalently because both reactions consume Ru5P and resolve the metabolic dead-end regardless of product identity. In contrast, the GLY-AUX design displayed a clear dependence on the oxygenase reaction for growth, with insensitivity to the carboxylation reaction (**Supplementary Fig. S1B, D**). Having established stoichiometrically that growth in the GLY-AUX background can be supported by Rubisco oxygenation alone, we next sought to study this coupling *in vivo*.

### A glycolate auxotroph couples growth to Rubisco oxygenation

*In vivo* Rubisco screening in *E. coli* requires co-expression with *prk*, since the Rubisco substrate RuBP is not naturally present in *E. coli*’s metabolic network (40). Because RuBP accumulation is detrimental, Rubisco’s activity alleviates this toxicity and thereby enables viable growth (41). However, when the system does not strictly depend on the Prk–Rubisco module, *prk* often acquires deleterious mutations (42).

Previous approaches to probe Rubisco’s activity *in vivo*, including pentose utilization deficiency (Δ*rpi*) (17, 37) and 3PG auxotrophy (43), cannot distinguish between carboxylation and oxygenation (**Extended Data Fig. 1B-E**). In contrast, coupling growth to 2PG metabolism provides a direct and exclusive readout of oxygenase activity (**Fig. 1A,B Supplementary Fig. S1**). 2PG is converted to glycolate by the constitutively expressed phosphoglycolate phosphatase gene (*gph*, EC 3.1.3.18) (44), and we recently developed *E. coli* glycolate auxotrophs capable of sensing this metabolite (38), providing a platform for linking Rubisco oxygenation to microbial fitness.

We therefore transformed the GLY-AUX2 (a derivative of the GLY-AUX strain (38) lacking the glycine cleavage system Δ*gcvTHP*; **Supplementary Fig. S2**), with the previously described pCBB plasmid carrying a constitutive *prk*-Rubisco module (*prk* from *Synechococcus elongatus* and *cbbM* from *R. rubrum*) (45) (**Fig. 1C**). Deletion of *gcvTHP* was motivated by the labeling profile previously observed, which suggested partial contribution of formate to glycine biosynthesis through reverse activity of the glycine cleavage system as consequence of heterologous expression of a formate tetrahydrofolate ligase (38, 46). As shown in **Fig. 1D**, strain GLY-AUX2 grew only when the pCBB plasmid was provided and the headspace contained ambient CO_2_ (0.04%). In contrast, growth was impaired at 10% CO_2_ (**Fig. 1D**). This suggests that at ambient CO_2_, Rubisco catalyzes oxygenation, generating 2PG that is further converted into glycolate, whereas at 10% CO_2_ the enzyme is saturated with CO_2_, allowing only carboxylation and thus preventing growth, thereby matching the *in silico* predictions.

To verify activity of the Prk-Rubisco module, we transformed pCBB into a Δ*rpi* strain and observed growth only in the presence of the plasmid (**Extended Data Fig. 1B-E**). Higher growth rates at 10% CO_2_ compared to ambient conditions likely reflect both the higher turnover rate of carboxylation versus oxygenation (47) and the additional metabolic cost of 2PG recycling when oxygenation occurs. Having confirmed that our platform selectively senses oxygenation through the activities encoded in the pCBB plasmid, we next developed a standardized expression platform to streamline cloning of diverse Prk and Rubisco variants.

### A modular, low-copy pSEVA-based plasmid architecture for tunable *prk*-Rubisco expression

We designed a new plasmid architecture for heterologous Prk–Rubisco module expression (**Fig. 1E**). The original pCBB plasmid is based on a p15A origin of replication and relies on a mutated, constitutively active P*_tet_* promoter (45), which largely restricts the host range and expression levels. To improve modularity, cloning flexibility and plasmid versatility, we introduced three key modifications (**Fig. 1E**). First, we substituted the mutated P*_tet_* promoter with a constitutive P*_j5_* variant (48, 49), which is functional in *E. coli* as well as in other Rubisco-containing bacteria such as *Cupriavidus necator.* Second, to standardize the cloning procedure, we transitioned to the Standard European Vector Architecture (SEVA) framework (50). Third, to improve the versatility of our plasmid setup, we adopted a broad-host-range, low-copy-number origin of replication (RK2), while replacing the resistance marker with a kanamycin resistance gene.

Because a highly active Prk is toxic to cells (42), we created a new transcription unit in which the translation initiation rate (TIR) value for *cfxP* from *Cupriavidus necator*, an alternative *prk* variant recently used for *in vivo* carboxylation screening (17), was set to ≈2000 a.u. We named this construct pCBB2. No mutations in *prk* were detected after heat-shock transformation of *E*. *coli* DH5α, and plasmid pCBB2 restored growth of strains Δ*rpi* and GLY-AUX2 (**Fig. 1F**). This TIR value did not lead to mutations in strain DH5α and enabled growth complementation of both Δ*rpi* and GLY-AUX2 (**Supplementary Fig. S3**). Notably, although the growth rate in *E*. *coli* GLY-AUX2 was slightly reduced (**Fig. 1G**), growth rates were comparable between pCBB and pCBB2 (0.09 ± 0.02 and 0.07 ± 0.02 h^-1^, respectively, two-tailed unpaired t-test p-value > 0.05). In contrast, growth rate decreased significantly in Δ*rpi* from pCBB (0.30 ± 0.01 h^-1^) to pCBB2 (0.13 ± 0.03 h^-1^), two-tailed unpaired t-test p-value < 0.0001 (**Supplementary Fig S4**), thereby establishing the activities borne by plasmid pCBB2 as a bottleneck for growth. Thus, the pCBB2 architecture represents a well-suited setup for testing different *prk* and Rubisco gene variants through growth-coupling.

### A diverse set of Prk variants is functionally interchangeable in pCBB2

To assess whether Prk identity limits oxygenation-dependent growth, we cloned eight variants spanning Alpha-, Beta-, and Gamma-proteobacteria, including both wild-type and codon-optimized sequences, into plasmid pCBB2*rbc4* (*prk1*–*prk8*, **Supplementary Data 1**). The reference construct *cfxP* from *C. necator*, hereafter *prk1*, was retained as the baseline. All variants were accommodated without introducing secondary mutations (**Fig. 2A, B**), confirming that Prk toxicity is mitigated in this architecture regardless of variant identity.

**Fig. 2.**
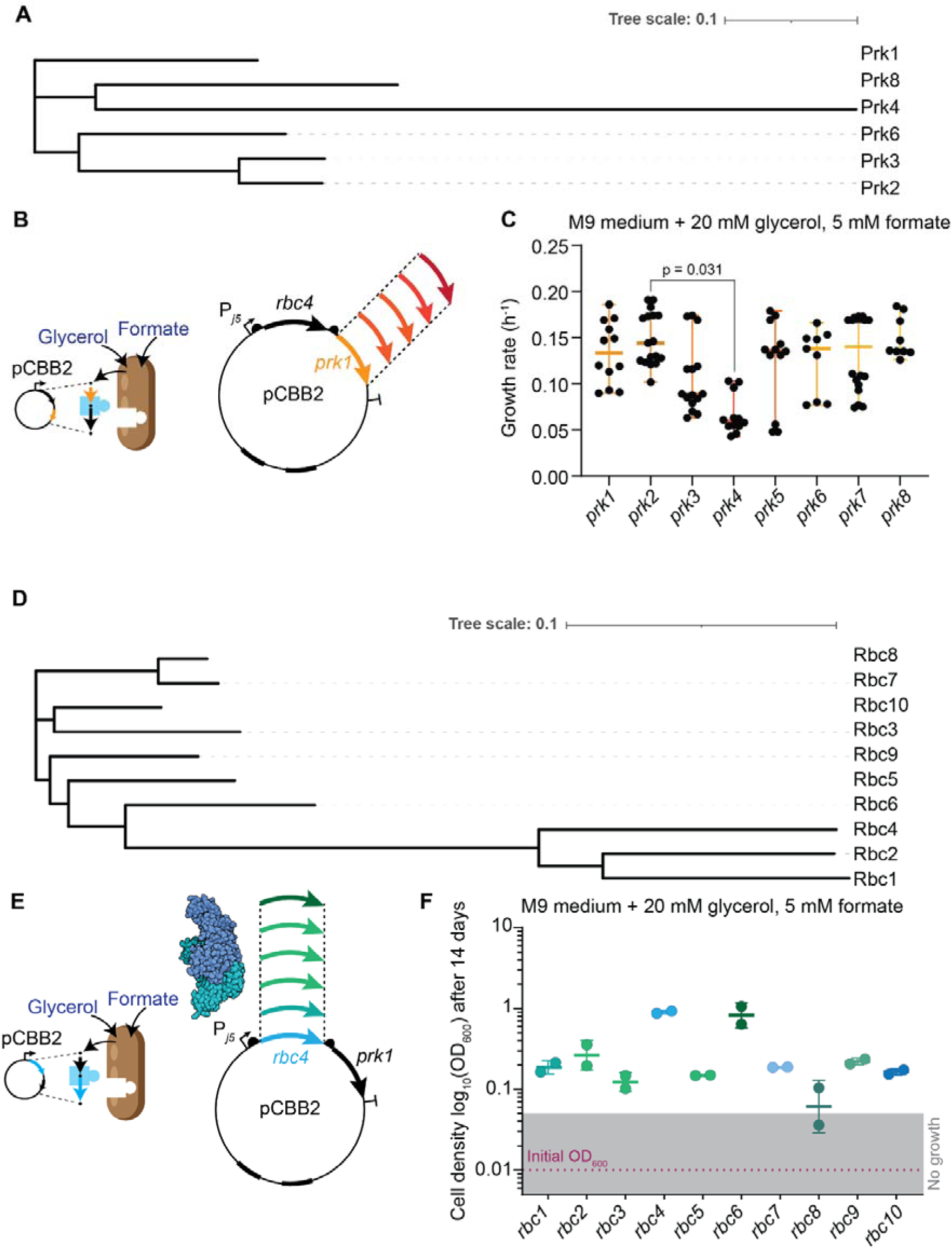
Phylogenetically diverse Prk and Form II Rubisco variants support oxygenation-dependent growth in the pCBB2 modular expression system. A) Phylogenetic tree of the eight Prk variants tested in this study. Variants span Alpha-, Beta-, and Gamma-proteobacteria classes within the Proteobacteria phylum.. Prk5 and Prk7 are codon-optimized versions of Prk4 and Prk6, respectively, and were therefore omitted from the phylogenetic tree. B) Schematic representation of the modular *prk* exchange strategy in pCBB2*rbc4*, and the resulting growth complementation through Prk–Rubisco flux under oxygenation-selective conditions. C) Growth rates of GLY-AUX2 strains harboring pCBB2*rbc4* variants carrying different *prk* genes at ambient CO_2_ (0.04%). Individual points represent the mean of technical triplicates from biologically independent experiments (n = 4–6); bars indicate the mean ± s.d. No significant differences were detected across variants (one-way ANOVA with Tukey’s multiple comparisons test), with the exception of *prk2* vs *prk4* (adjusted p = 0.031, indicated in figure). D) Phylogenetic tree of the ten Form II Rubisco variants tested in this study, spanning Alpha-, Beta-, Gamma-, and Zeta-proteobacteria classes. E) Schematic representation of the modular *rbc* exchange strategy in pCBB2*prk1*, and the resulting growth complementation through oxygenase activity. F) Final OD_600_ values after 14 days of incubation in sealed baffled shake flasks under oxygenation-selective conditions (ambient CO2). Individual points represent independent biological replicates (n=2); bars indicate the mean ± s.d. *rbc4* and *rbc6* reached the highest final OD_600_ values across the ten Form II Rubisco variants tested in GLY-AUX2, consistent with their superior performance in plate-reader cultivations.

Growth rates were comparable across all constructs, with no variant significantly outperforming Prk1 in GLY-AUX2 (**Fig. 2C, Supplementary Fig. S6**), except for a marginal difference between *prk2* and *prk4* (one-way ANOVA; adjusted p = 0.031). Oxygenation-dependent growth of strain GLY-AUX2 is therefore largely insensitive to Prk sequence and codon optimization, validating pCBB2 as a robust expression platform for diverse Prk variants. We retained *prk1* for all subsequent experiments.

### Phylogenetically diverse Form II Rubiscos support oxygenation-dependent growth

Next, we replaced the Rubisco gene from *R. rubrum* (*rbc4*) in pCBB2 with nine additional Form II Rubisco variants selected for high *k_cat_* values *in vitro* (51) and spanning Alpha-, Beta-, Gamma-, and Zeta-proteobacteria (**Fig. 2D, E, Supplementary Fig. S7, Supplementary Data 1**). Form II Rubiscos were prioritized for their compatibility with native *E. coli* chaperones (52) and their relatively high oxygenation-to-carboxylation ratio compared to other Rubisco forms (53).

We first checked whether the cloned Prk–Rubisco modules were catalytically functional by assessing growth complementation by the same pCBB2 variants transformed in the Δ*rpi* background, selecting for carboxylation at 10% CO_2_. Eight out of ten variants reached stationary phase within 150 h (**Supplementary Fig. S8**), confirming that most modules actively support metabolic flux through the carboxylation reaction.

We then assessed growth complementation of GLY-AUX2 transformed with each pCBB2 variant under ambient pCO_2_ (0.0004 atm or 0.04% CO_2_) in plate-reader cultivations. Despite confirmed module functionality, only *rbc4* and *rbc6* from *Sulfurivirga caldicuralii* supported growth (**Supplementary Fig. S9**). We hypothesized that ineffective flux through Prk–Rubisco, possibly caused by imbalanced protein stoichiometry, reduced oxygenation activity below the threshold required for growth. Unlike Δ*rpi*, GLY-AUX2 growth depends strictly on oxygenation-derived products with no alternative metabolic route, and carboxylation and oxygenation compete under ambient CO_2_, together imposing a higher demand on Rubisco oxygenation flux.

Consistent with this interpretation, analysis of predicted TIR values revealed that when *rbc4* was held constant and *prk* variants were substituted **(Fig. 2A-C**), Rubisco-to-Prk TIR ratios remained within 1–10 arbitrary units across nearly all constructs (**Supplementary Table S2**). By contrast, fixing *prk1* while varying the Rubisco introduced substantially greater variability, with ratios spanning from 2 to over 200 (**Supplementary Table S3**). Such stoichiometric imbalance could thereby limit effective oxygenation flux in strain GLY-AUX2, where the metabolic demand on Rubisco-derived products is high.

We therefore reasoned that oxygen availability in the cultures could compound this limitation and performed growth complementation experiments in sealed baffled shake flasks, where evaporation is prevented and any OD_600_ increase above background strictly requires sustained Rubisco-mediated oxygenation. Accordingly, we observed significant turbidity increases for nearly all tested variants, with *rbc4* and *rbc6* mediating the highest final OD_600_ values (**Fig. 2F**), consistent with plate-reader results. Growth was modest when the selection strain was transformed with most variants (OD_600_ ≤ 0.4), except for *rbc4* and *rbc6* (OD_600_ > 1.0). Yet, the data demonstrates that oxygenation-dependent complementation is achievable across phylogenetically diverse Form II Rubiscos although the overall performance ceiling of the GLY-AUX2 background nonetheless remained low. Therefore, we next applied ALE to improve module compatibility and functional output.

### Adaptive laboratory evolution localizes the oxygenation bottleneck to the Prk–Rubisco module

The modest oxygenation-dependent growth rates achieved with pCBB2*rbc4prk1* in GLY-AUX2 represented an opportunity to apply the selective pressure of continuous cultivation to improve module performance *in vivo*. We therefore subjected GLY-AUX2 harboring pCBB2rbc4prk1 to ALE in turbidostatic mode operated in four independent reactors (R1–R4) for five weeks (**Fig. 3A, B, Extended Data Fig. 2A**). A consistent decrease in time between dilutions confirmed progressive improvement in population growth rate (**Extended Data Fig. 2B, C**). Two parallel turbidostat runs under relaxed conditions (R5–R6, strain GLY-AUX2 carrying the empty pSEVA221 vector) served as controls to filter mutations arising from genetic drift or platform adaptation.

**Fig. 3.**
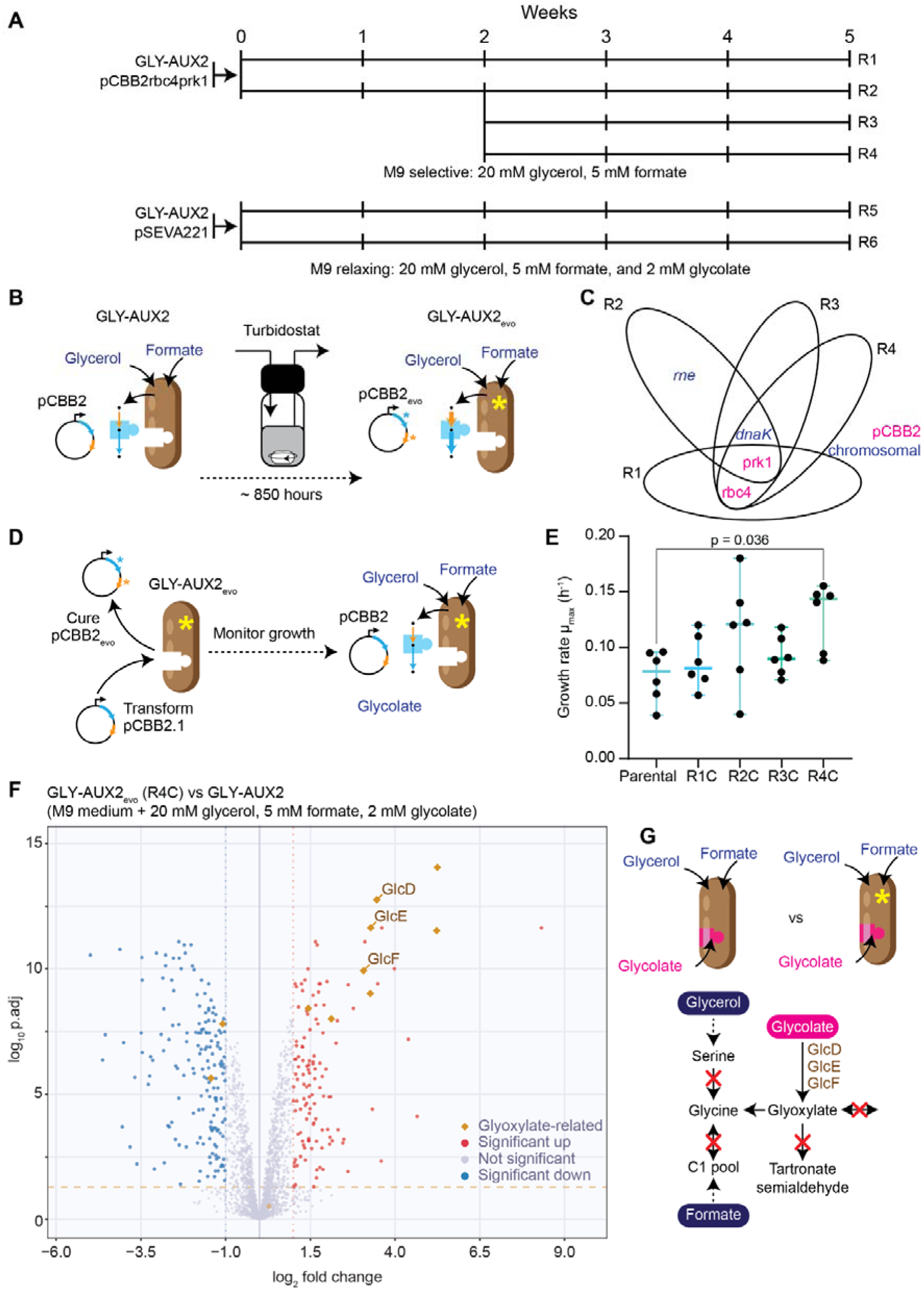
Adaptive laboratory evolution identifies mutations in the Prk–Rubisco module as the primary driver of improved oxygenation-dependent growth. A) Schematic of the turbidostat setup showing the biological content and evolutionary trajectories of reactors R1–R4 (selective conditions) and R5–R6 (relaxed conditions, controls) over five weeks of continuous cultivation. B) Schematic of the turbidostat ALE strategy, illustrating the adaptive mutations under continuous oxygenation-selective pressure. C) Venn diagram of mutations identified across R1–R4 populations (variant frequency ≥70%); chromosomal mutations in navy blue, plasmid-borne mutations in magenta; overlapping regions indicate shared mutations. D) Schematic of the plasmid curing and retransformation strategy: GLY-AUX2_evo_ isolates from R1–R4 were cured of evolved pCBB2 and retransformed with naïve pCBB2r*bc4prk1* to isolate the contribution of chromosomal mutations to the evolved phenotype. E) Growth rate comparison between GLY-AUX2_evo_ isolates from R1–R4 retransformed with naïve pCBB2*rbc4prk1* compared to parental GLY-AUX2 at ambient CO_2_. Each dot represents technical replicates from biological duplicates. Data are mean ± s.d. (n=2 biologically independent experiments); individual points represent technical replicates. The R4-derived isolate showed a modest but significant improvement (Tukey’s multiple comparisons test, adjusted p=0.036); remaining isolates were not significantly different from the parental strain. F) Volcano plot from proteomic analysis of a representative GLY-AUX2evo clone from R4, showing differentially expressed proteins relative to the parental strain. G) Schematic illustrating upregulation of GlcDEF in evolved isolates. Although increased GlcDEF abundance may improve processing of oxygenation-derived glycolate, chromosomal adaptations are not the primary driver of the evolved phenotype.

Whole-genome sequencing of evolved populations revealed no chromosomal mutations that could be traced to glycolate or glyoxylate metabolism (**Fig. 3C, Supplementary Data 2**), localizing the adaptive bottleneck to the plasmid-borne Prk–Rubisco module. Plasmid sequencing identified two recurrently selected mutations: M115I in *rbc4* (hereafter *rbc4*\*) and N216T in *prk1* (hereafter *prk1*\*) (**Fig. 3C, Table S4**). In R2, *prk1\** arose first with *rbc4\** emerging only after an additional four weeks, suggesting sequential rather than simultaneous selection (**Table S5**).

To completely rule out the effect of the chromosomal mutations, we isolated single GLY-AUX2_evo_ clones from R1-R4 were cured of the evolved plasmid and retransformed with the naïve (non-evolved) pCBB2*rbc4prk1* plasmid (**Fig. 3D**). Growth rates were largely comparable to the parental strain across isolates, with the exception of the R4-derived clone, which showed a modest but statistically significant improvement (Tukey’s multiple comparisons test, adjusted p = 0.036; **Fig. 3E, Supplementary Fig. S10**). Proteomic analysis of these isolates revealed a consistent upregulation of proteins involved in glyoxylate and dicarboxylate metabolism in all four clones (**Fig. 3F, G; Supplementary Files 2 and 3**). Taken together, these results indicate that chromosomal adaptations (most notably upregulation of glyoxylate metabolism) contribute marginally to the evolved phenotype in at least one isolate but are unlikely to account for the substantial fitness gains observed during ALE. We therefore focused on the plasmid-borne mutations *prk1*\* and *rbc4*\* as the primary drivers of improved oxygenation-dependent growth.

### N216T substitution in *prk1* enhances Prk-Rubisco flux independent of reaction selectivity

Next, we assessed the impact of the mutation at the Prk level. To this end, the N216T substitution was reverse engineered into *prk1* in pCBB2rbc4prk1* (**Fig. 4A**) and tested in the naïve GLY-AUX2 strain under ambient pCO_2_ conditions (**Fig. 4B**). The resulting strain exhibited a 2.7-fold increase in growth rate compared to the control strain harboring the original pCBB2*rbc4prk1* plasmid, increasing from 0.12 ± 0.02 h^-1^ to 0.32 ± 0.01 h^-1^ (unpaired two-tailed t-test for all this section, p-value = 0.00001), respectively (**Fig. 4C, D**). Additionally, the lag phase for growth was shortened as well, from 15.3 ± 2.1 to 4.4 ± 0.6 h, respectively (p-value = 0.001). This result indicates an improvement in oxygenation-dependent flux associated with the Prk1 variant. Notably, as will be described in the following section, introduction of Prk1* resulted in observable oxygenation-dependent growth of Rbc4 also at 0.5% CO_2_, representing an approximately 10-fold increase in pCO_2_ compared to ambient conditions.

**Fig. 4.**
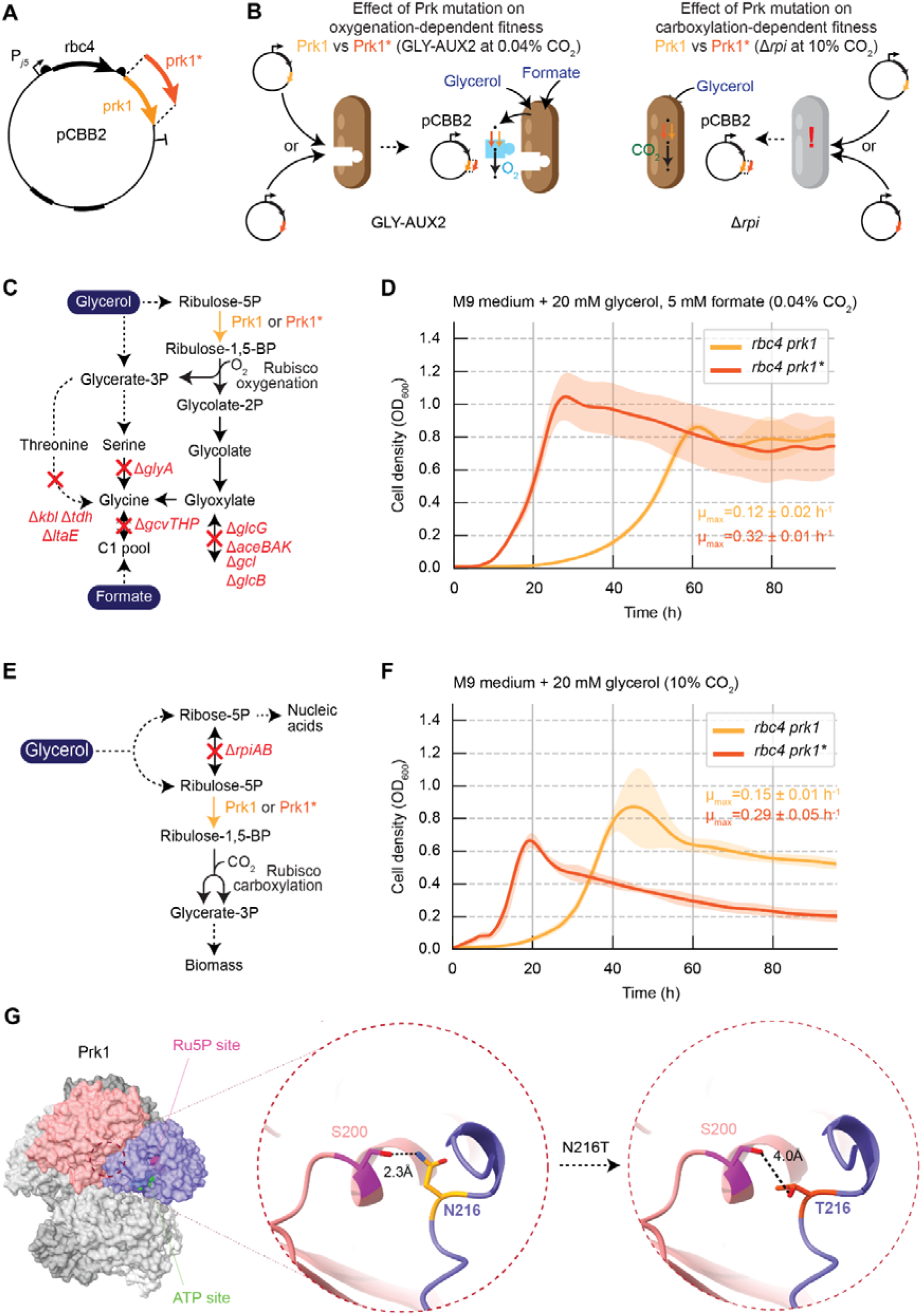
The N216T substitution in Prk1 enhances Prk–Rubisco flux independently of reaction selectivity. A) Schematic of the retroengineering strategy: the N216T substitution (*prk1\**) was introduced into naïve pCBB2*rbc4prk1* to assess its contribution to the evolved phenotype independently of other ALE-derived mutations. B) Experimental design overview: the effect of *prk1\** was evaluated in two complementary strain backgrounds — GLY-AUX2, reporting on oxygenase activity, and Δ*rpi*, reporting on carboxylation activity — using the same Rubisco variant (*rbc4*) in both contexts. C) Schematic of GLY-AUX2 under oxygenation-selective conditions, highlighting the metabolic flux difference between strains harboring *prk1* and *prk1\**. D) Growth profiles of GLY-AUX2 strains harboring pCBB2*rbc4prk1* or pCBB2*rbc4prk1\**, measured at ambient CO_2_ (0.04%). The prk1* variant showed a 2.7-fold increase in growth rate relative to prk1 (p-value < 0.0001). E) Schematic of Δ*rpi* under carboxylation-selective conditions at 10% CO_2_, highlighting the metabolic flux difference between strains harboring *prk1* and *prk1\**. F) Growth profiles of Δ*rpi* strains harboring pCBB2*rbc4prk1* or pCBB2*rbc4prk1\**, measured at 10% CO_2_. The *prk1\** variant showed an approximately 2-fold increase in growth rate relative to *prk1* (p-value < 0.01). G) AlphaFold-based structural model of the Prk1 octamer, showing the location of N216 at a subunit interface, approximately 20–21 Å from the RuBP-and ATP-binding sites. The N216T substitution is predicted to alter the geometry of the intersubunit contact with S200 from a neighboring protomer, potentially modulating oligomerization dynamics and catalytic turnover.

We reasoned that the *prk1*\* mutation could debottleneck the Prk-Rubisco module also when paired with other Rubisco variants. Therefore, we introduced *prk1*\* in two pCBB2 plasmids containing *rbc2* and *rbc3*, respectively. Then, we repeated the shake flask cultivations and determined cell turbidity after 14 days of incubation. Although we could determine a significant 4.5-fold increase in final OD_600_ value only for the *rbc3* construct (from 0.13 ± 0.03 to 0.54 ± 0.04, p-value = 0.0075) (**Supplementary Fig. S11**), this result confirms that the nonsynonymous N216T substitution in Prk1 can increase the Prk-Rubisco flux also in other Rubiscos besides Rbc4.

We next evaluated the effect of the mutation in the context of carboxylation-dependent growth using the Δ*rpi* strain at 10% CO_2_ with Rbc4 (**Fig. 4E**). Here again, expression of Prk1* enhanced growth relative to the original plasmid, resulting in ≈2-fold increase in growth rate from 0.15 ± 0.01 to 0.29 ± 0.05 h^-1^ (p-value = 0.008) (**Fig. 4F**). Also in this case, the lag phase reduced as an effect of *prk1\** from 6.8 ± 2.1 to 0.6 ± 1.0 hours, respectively (p-value = 0.006).

Collectively, these results demonstrate that the nonsynonymous N216T substitution in *prk1* enhances flux through Prk and consequently provides RuBP at a higher rate for the Prk-Rubisco module, independently of whether growth is supported by Rubisco oxygenation or carboxylation. Because the same direction of improvement appears under both selection conditions, Prk1* is most parsimoniously interpreted as relieving the RuBP-supply bottleneck rather than altering Rubisco selectivity; a reaction-agnostic, flux-level improvement of the Prk–Rubisco module. This inference is supported by the orthogonal readout from the two selection strains, which jointly distinguish module-level flux effects from Rubisco-selectivity effects. To gain insight into the mechanistic basis of the ALE-derived mutation, we examined the structural context of the substituted residue in Prk. Mapping the mutation onto the Prk1 AlphaFold-structure revealed that N216 is located on the solvent-exposed surface of the protomer and lies approximately 20–21 Å from the RuBP- and ATP-binding sites, suggesting that its effects are unlikely to arise from direct alterations of the catalytic center. Given that Prk1 has been reported to assemble as an octamer (54, 55), we generated an AlphaFold-based octameric model to assess potential intersubunit interactions (**Fig. 4G**). In this model, N216 resides at a subunit interface and is predicted to form a hydrogen bond with S200 from a neighboring protomer. Replacement of asparagine with threonine shortens the side chain and replaces the terminal amide with a hydroxyl group, potentially altering the geometry or stability of this interaction while preserving its polarity. Although the biochemical consequences of this substitution remain to be determined, the structural model raises the possibility that the mutation could influence conformational coupling between subunits or oligomerization dynamics consistent with increased catalytic turnover, thereby modulating Prk1 activity and the supply of RuBP to the Rubisco-mediated reaction. We therefore retained this enhanced Prk-Rubisco flux background as the genetic context for assessing the effect of the Rubisco mutant.

### Rbc4^M115I^ shifts the *in vivo* balance toward oxygenation

Next, we introduced the M115I substitution in *rbc4* within the pCBB2*rbc4prk1\** plasmid and evaluated its impact on oxygenation-dependent growth in the GLY-AUX2 system (**Fig. 5A-C**). At 0.04% CO_2_, the plasmid harboring Rbc4* supported only a modest increase in growth rate and final biomass relative to the original pCBB2rbc4prk1 plasmid (unpaired two-tailed t-test, p-value = 0.4), with lag phase of 30.5 ± 4.1 hours. In contrast, the strain producing Prk1* and the original Rbc4 displayed the highest growth performance under these conditions (**Supplementary Fig. S12**).

**Fig. 5.**
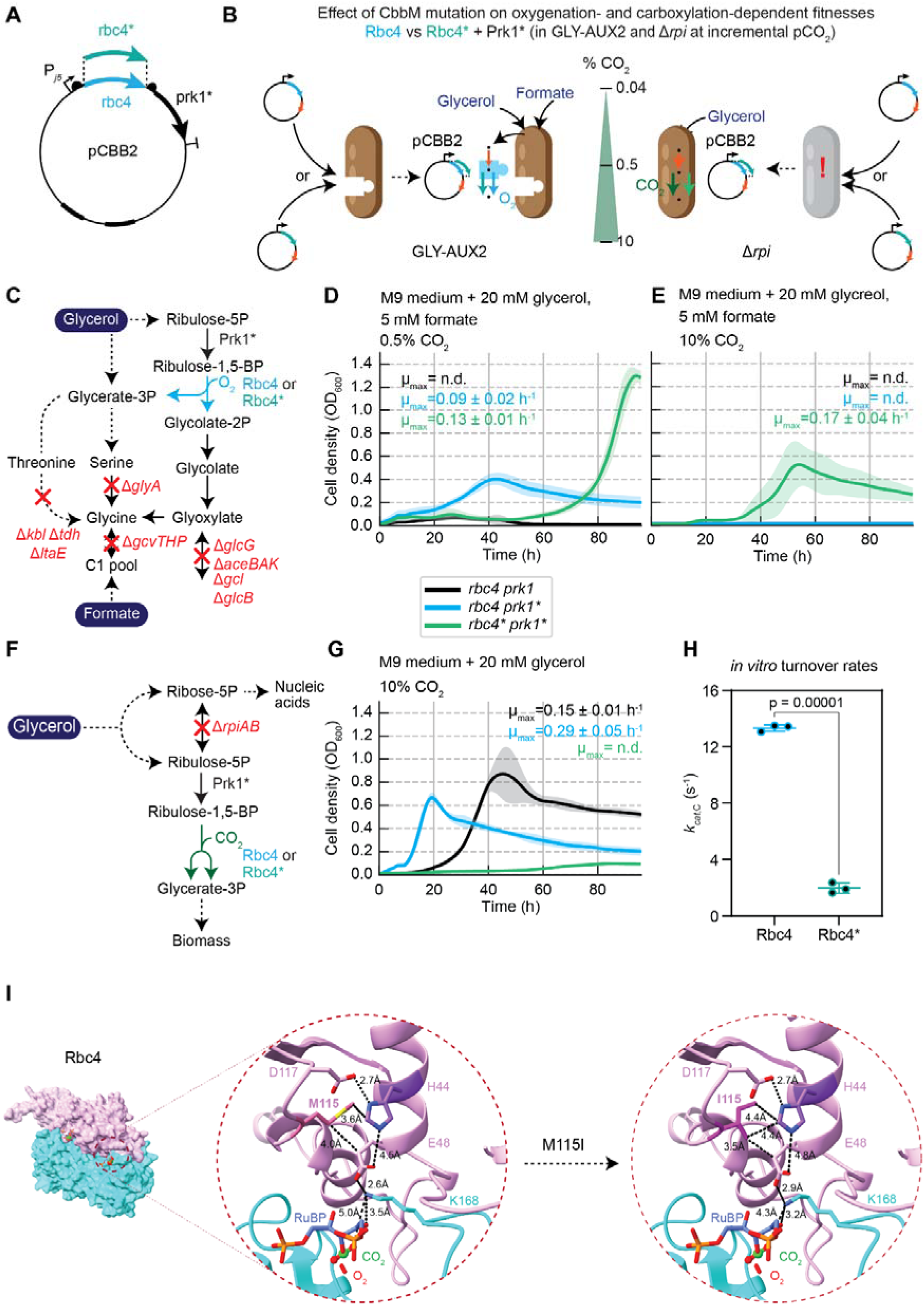
The M115I substitution in Rbc4 shifts the in vivo carboxylation/oxygenation balance toward oxygenation. A) Schematic of the retroengineering strategy: the M115I substitution (*rbc4*\*) was introduced into naïve pCBB2*rbc4prk1\** to assess its contribution to the evolved phenotype independently of other ALE-derived mutations. B) Experimental design overview: the effect of *rbc4*\* was evaluated in GLY-AUX2 and Δ*rpi* backgrounds across a range of CO_2_ concentrations (0.04%, 0.5%, and 10% CO2), using *prk1\** as the fixed Prk variant. C) Schematic of GLY-AUX2 under oxygenation-selective conditions, highlighting the metabolic flux difference between strains harboring *rbc4* and *rbc4*\*. D) Growth profiles of GLY-AUX2 strains harboring pCBB2*rbc4prk1*, pCBB2*rbc4prk1\**, and pCBB2*rbc4*prk1\**, measured at 0.5% CO_2_. The *rbc4prk1* module showed a 1.4-fold higher growth rate (p-value < 0.05) and 3-fold higher final biomass concentration (p-value < 0.0001) compared to *rbc4prk1\**. The curves shown here represent one biological replicate measured in technical quadruplicate. Shaded areas represent standard error with 95% confidence interval. Experiments were performed in biological triplicate. E) Growth profiles of the same three constructs measured at 10% CO_2_. Detectable growth was observed exclusively in the strain harboring the *rbc4prk1* module, reaching a growth rate of 0.17 ± 0.04 h⁻¹ and a maximal OD of 0.54 ± 0.20. Curves represent one biological replicate measured in technical quadruplicate. Shaded areas represent standard error with 95% confidence interval. Experiments were performed in biological triplicate. F) Schematic of Δ*rpi* under carboxylation-selective conditions at 10% CO_2_, highlighting the metabolic flux difference between strains harboring *rbc4* and *rbc4\**. G) Growth profiles of Δ*rpi* strains harboring pCBB2*rbc4prk1*, pCBB2*rbc4prk1\**, and pCBB2*rbc4prk1* at 10% CO_2_. Production of Rbc4* together with Prk1* severely impaired growth, with virtually no detectable cell division. Curves represent one biological replicate measured in technical quadruplicate. Shaded areas represent standard error with 95% confidence interval. Experiments were performed in biological triplicate. H) *In vitro* comparison of carboxylation turnover rates (*k_cat,C_*) between Rbc4 and Rbc4*. The M115I substitution results in a 6-fold reduction in *k_cat,C_*, from 14.2 ± 0.2 s⁻¹ to 2.1 ± 0.4 s⁻¹ (mean ± standard deviation, statistical comparisons of technical triplicates by two-tailed unpaired t-test; exact p-values are indicated). I) AlphaFold-based structural model of Rbc4, showing the network of residues proximal to M115, including H44, E48, and K168. The M115I substitution is predicted to alter hydrophobic contacts within this network, potentially constraining the geometry of the catalytic pocket and compromising carboxylation-specific turnover through indirect effects on active-site organization.

The advantage conferred by the M115I substitution in Rbc4 became more apparent at pCO_2_ conditions above ambient CO_2_ saturation. In fact, in a turbidostat cultivation, it is likely that the local pCO_2_ inside the cells is higher due to the constant catabolism of the heterotrophic substrates (mainly glycerol). As mentioned in the previous section, introduction of *prk1*\* allowed us to demonstrate oxygenation dependent growth for *rbc4* at 0.5% CO_2_. Under this pCO_2_ value, the *rbc4*prk1\** module showed a 1.4-fold higher growth-rate than the *rbc4prk1\** module (0.09 ± 0.02 vs 0.13 ± 0.01 h^-1^, respectively, p-value = 0.028) and reached a 3-fold higher final biomass concentration (OD_600_ of 1.35 ± 0.02 versus 0.41 ± 0.05, respectively, p-value = 0.000001), albeit with a lag phase of 50.4 ± 1.3 h (**Fig. 5D**). The extended lag phase notwithstanding, the superior growth rate and final biomass of the *rbc4prk1* module represent a clear fitness advantage in a turbidostat selection context, where sustained exponential growth determines competitive outcome. In contrast, the parental pCBB2*rbc4prk1* construct did not support any growth in strain GLY-AUX2 incubated at 0.5% CO_2_ and above (**Fig. 5D**).

The fitness advantage of the *rbc4\** mutation became even more pronounced at 10% CO_2_, where detectable growth was observed exclusively in the strain harboring the *rbc4*prk1\** module, reaching a growth rate of 0.17 ± 0.04 h^-1^ and a maximal OD_600_ of 0.54 ± 0.20, and with a lag phase of 22.3 ± 2.3 h (**Fig. 5E**). Together, these results indicate that the M115I substitution enhances oxygenation-dependent growth fitness, particularly under elevated CO_2_ conditions.

To determine whether this improvement occurred at the expense of carboxylation, we tested the same constructs in the Δ*rpi* background at 10% CO_2_ (**Fig. 5F**). Under these conditions, growth depends exclusively on carboxylation flux. Therefore, any growth impairment reflects reduced carboxylation activity rather than increased oxygenation. Indeed, production of Rbc4* together with Prk1* severely impaired growth, with virtually no detectable cell division (**Fig. 5G**), indicating that the M115I substitution shifts the *in vivo* catalytic balance of Rbc4 toward oxygenation at the expense of carboxylation.

*In vitro* kinetic measurements confirmed and quantified this loss of carboxylation activity, revealing a 6-fold reduction in *k_cat,C_*, decreasing from 14.2 ± 0.2 s⁻¹ for the wild-type enzyme to 2.1 ± 0.4 s⁻¹ for Rbc4* (p = 0.00001) (**Fig. 5H**). To rationalize this effect structurally, we examined experimentally determined structures of Rubisco from *R. rubrum* and *Galdieria sulphuraria*. The catalytic substrates (RuBP, CO_2_ and O_2_) are predicted to bind approximately 10–13 Å from residue 115, which is therefore unlikely to participate directly in substrate binding. Nevertheless, structural modelling suggested subtle local rearrangements in a network of residues involving H44, E48 and K168, amino acids that are three-dimensionally proximal to position 115 despite being distant in the primary sequence. Replacement of methionine with isoleucine alters the geometry of hydrophobic contacts with H44 and, more importantly, is predicted to affect packing interactions involving E48, which is positioned near the RuBP-binding residue K168 (**Fig. 5I**). These structural features suggest that M115I compromises carboxylation-specific turnover through indirect effects on active-site organization rather than direct substrate interactions. Intriguingly, previous directed-evolution studies (47) align with our hypothesis as they identified functional mutations at H44 and D117, two residues spatially flanking M115, that reduced the relative efficiency of carboxylation over oxygenation, further corroborating the proposed relevance of this region for Rubisco activity.

Whether this reflects a true gain in oxygenase catalytic activity or a relative reweighting of *in vivo* flux driven by impaired carboxylation will require direct measurement of *k*_cat,O_ on the purified variant; either outcome is consistent with a single-substitution shift in the *in vivo* catalytic balance of Form II Rubisco that is accessible to growth-coupled selection.

## Discussion

Rubisco’s oxygenase activity has been characterized biochemically for decades but has never been accessed as a direct selection trigger *in vivo*. Two previously reported *E. coli* platforms assay Rubisco activity by coupling RuBP turnover to growth — the Δ*gapA* strain through a Prk–Rubisco shunt around a glycolytic deficiency (42, 47, 56–58), and Δ*rpi* by detoxifying Ru5P accumulation through a plasmid-borne Prk–Rubisco module (17, 37, 42, 47) — but neither resolves the carboxylation–oxygenation partition that defines Rubisco’s central catalytic trade-off, because growth depends on RuBP consumption regardless of whether the products are 3PG or 2PG. Here, we developed a complementary platform based on glycolate auxotrophy. Strain GLY-AUX2 couples 2PG formation to glycine biosynthesis through an engineered glycolate auxotrophy (38), with deletion of *gcvTHP* blocking the reverse, CO_2_-fixing direction of the glycine cleavage system that would otherwise supply glycine independently of Rubisco oxygenation at elevated pCO_2_. To our knowledge, no previous strain has coupled microbial growth specifically to Rubisco oxygenation, enabling both screening and continuous selection for an activity that has been largely inaccessible to forward-genetic approaches.

Pairing the modification in strain GLY-AUX2 with the Δ*rpi* deletion allowed us to read out oxygenation and carboxylation fluxes, respectively, from the same Rubisco variant in two complementary genetic backgrounds, and this dual readout proved diagnostic for the ALE-derived substitutions Rbc4* and Prk1*. Rbc4* showed a reduction in growth for Δ*rpi*, corroborated by a 6-fold reduction in *k_cat,C_ in vitro* yet enhanced oxygenation-dependent growth in GLY-AUX2; this is direct *in vivo* evidence that a single amino acid substitution can shift the carboxylation/oxygenation balance of a Form II Rubisco. M115 lies adjacent to D117, a residue previously implicated in modulating *S*_C/O_ through reductions in *k_cat,C_* and *k_cat,C_*/*K_C_*. Structural modelling provides a plausible mechanistic basis for this phenotype. The M115I substitution alters a network of hydrophobic contacts involving H44 and E48, both residues proximal to the active site of *R. rubrum* Rubisco, in a way that repositions first E48 and consequently K168 (already forming a hydrogen bond with the RuBP substrate), likely modifying the native geometry of the catalytic pocket. This rearrangement is consistent with a reduction in RuBP accommodation without proportional loss of oxygenase activity, providing a structural rationale for the observed reduction in *k_cat,C_*. In the absence of direct *S*_C/O_ measurements and an experimentally determined structure of the variant in complex with the substrates, this interpretation remains tentative. Yet, the *in vivo* and *in vitro* phenotypes establish the carboxylation–oxygenation trade-off independently of the specific mechanism of action of the ALE-derived mutation. Prk1*, in contrast, increased growth under both carboxylation- and oxygenation-selective conditions and is most parsimoniously explained as a non-selectivity-altering improvement in RuBP supply, which becomes available for Rubisco. This illustrates that the dual readout also distinguishes selectivity-shifting mutations from generic fitness improvements.

Four features of the platform deserve to be framed as design insights rather than incidental limitations. First, oxygenation-dependent growth in GLY-AUX2 required baffled shake flasks to be detectable across the full variant panel; in standard plate-reader cultivation, only two of ten Rubiscos supported measurable growth, while carboxylation screening in the Δ*rpi* strain at 10% CO_2_ was robust (8/10 variants). This O_2_-sensitivity is mechanistically expected: the assay is selective for oxygenation precisely because growth tracks RuBP to 2PG flux; therefore, dissolved O_2_ is a tunable selection variable for future high-throughput setups. Second, growth fitness in GLY-AUX2 integrates Rubisco activity with RuBP supply, intracellular CO_2_/O_2_ partitioning, enzyme expression and folding, and downstream glyoxylate metabolism. Several built-in controls anchor the interpretation of this system-level readout: the strain does not grow without the Prk–Rubisco module (**Extended Data Fig. 1 F-G**), (unless evolved) does not grow at elevated CO_2_ where Rubisco oxygenation is suppressed (**Fig. 1D**), and is stoichiometrically predicted by modeling to depend exclusively on the oxygenation flux (**Fig. 1A, B, Extended Data Fig. 1A, Supplementary Fig. S1**). Attribution of fitness differences across variants is further anchored by the orthogonal Δ*rpi* readout, which distinguishes selectivity-altering from generic flux-altering mutations — as illustrated by the Rbc4* and Prk1* variants, respectively — and by *in vitro* characterization of selected variants, as for the *k_cat,_*_C_ measurement on Rbc1*. Third, the pCBB2 expression architecture was tuned against a single Prk–Rubisco pair (rbc4 and prk1), and the resulting RBS strengths are unlikely to be optimal across the full range of screened variants; reaching plate-reader-detectable oxygenation growth for each variant will likely require a per-variant round of RBS retuning, which the modular design of pCBB2 supports. Fourth, the two sensor strains impose different selective demands on the heterologous module. In Δ*rpi*, carboxylation needs only supplying a small fraction of total carbon flux in order to complement growth (42), whereas in GLY-AUX2 the oxygenation-derived glycine must satisfy a larger proportion of cellular nitrogen and one-carbon demands, making GLY-AUX2 a more stringent selection platform in terms of the absolute Rubisco flux required for detectable growth. This higher activity threshold is a disadvantage for variants resulting in poor flux through the module for several reasons e.g., low protein production, solubility, low *k_cat_* in the bottleneck enzyme, or high *K_M_*. The difference in selective stringency between the two strains is worth designing around, and points to the value of their pairing as demonstrated here: using Δ*rpi* to confirm active flux through pCBB2 variants that fail to complement GLY-AUX2 disambiguates expression or folding failures from genuine insufficiency of oxygenation activity.

Our ALE campaign nominally selected at ambient air, but glycerol oxidation elevated dissolved CO_2_ in the bioreactor and produced a selection regime in the liquid phase closer to the elevated-pCO_2_ conditions under which Rbc4* subsequently outperformed wild-type. This is a reminder that the effective selection pressure in continuous cultivation depends on dissolved pCO_2_/pO_2_, not just headspace composition.

The ability to select directly on Rubisco’s oxygenase activity opens lines of investigation that have been difficult to pursue with carboxylation-coupled assays alone. Oxygenation is the less-studied of Rubisco’s two reactions: it is slow, thermodynamically wasteful, and historically assayed only *in vitro* under conditions that incompletely recapitulate the cellular environment. The GLY-AUX2 strain places oxygenation under direct growth selection, making fitness a proxy for this reaction in a living cell. This enables systematic mutational dissection of the sequence determinants of oxygenase activity across Form II and, in principle, Form I phylogenies, a landscape that *in vitro* surveys have only partially resolved (6, 7, 17, 22, 26, 51, 59), and one that is particularly relevant to understanding how Rubisco has evolved under varying atmospheric O_2_ and CO_2_ regimes (8, 26, 60–62). Because the assay is bacterial and growth-coupled, it scales to library sizes inaccessible to *in vitro* kinetics, making the molecular basis of oxygenation experimentally tractable at a resolution not previously available. By the same token, the same growth coupling that we use here to *select for* oxygenation could be in principle inverted to select *against* it, providing a forward-genetic route toward Rubisco variants with suppressed oxygenase activity — a long-standing goal of enzymology, synthetic biology, plant sciences, and crop engineering (4, 14, 63, 64). Pursuing these directions will require pairing GLY-AUX2 with higher-throughput aerated cultivation and with structural and computational analyses that connect *in vivo* fitness to active-site geometry, but the foundation is now in place.

## Data availability

Data supporting the findings of this work are available within the paper and its Supplementary Data Files. A reporting summary for this Article is available as a Supplementary Information file. Source data are provided in this paper. All strains presented in the manuscript can be obtained for academic research from the corresponding author upon request. Source data are provided in this paper.

## Code availability

All scripts, data files, and result plots can be found on our GitLab repository: https://github.com/MathiasWagner-99/RuBisCO-Oxygenation-Data-analysis/tree/main.

## Materials and methods

### Modelling of Rubisco oxygenation activity

In order to calculate phenotypic phase planes (rubisco carboxylation flux vs. biomass production) for two the rubisco-growth-coupled mutants, we adjusted the recently published *i*CH360 *E. coli* model (39) by adding the following relevant reactions (along with their reactants, when necessary): phosphoribulokinase (PRK), rubisco carboxylation and rubisco oxygenation (RBC and RBC_OX), the two Entner-Doudoroff reactions (EDD and EDA), phosphoglycolate phosphatase (PGLYCP), glycolate oxidase (GLYCTO1), and glycine-oxaloacetate transaminase (GLYOAT).

We then used the COBRApy package (65) to calculate the phenotypic phase planes for all combinations of the following parameters: (i) carbon source - glycerol, xylose, or glucose, (i) mutant - Δ*RPI*Δ*EDD*Δ*EDA* (i.e. Δrpi in short) or Δ*ICL*Δ*GHMT2r* (i.e. the GLY-AUX2), (iii) target reaction - rubisco carboxylation or rubisco oxygenation. When plotting the phase planes for rubisco carboxylation, the oxygenation reaction was completely shut off (set both bounds to 0) - and vice versa (**Figure S1**).

For the rest of the analysis we focused only on the GLY-AUX2 mutant with glycerol as the carbon source. In order to explore the effect of mixing carboxylation and oxygenation at different fixed ratios (defined by φ), we added a linear constraint on a combination of carboxylation and oxygenation fluxes: v(RBC) - ratio*v(RBC_OX) = 0. The scalar value “ratio” was set as 10^φ^, where φ acquires different values between −4 and 4. In **Fig. 1B**, we combined the ratio constraint with an upper bound on the sum of both rubisco fluxes (set to 1 μmol/min/gCDW). We then maximized growth of the GLY-AUX2 mutant growing on glycerol (at each φ). Since carboxylation is counter-productive in this mutant, the function is monotonically increasing with φ (i.e. more oxygenation is better).

For generating the phenotypic phase planes for different values of φ (**Extended Data Fig. 1A**), each phase plane is computed by sweeping the flux in RBC, while again pinning the flux ratio between v(RBC_OX) and v(RBC) to 10^φ^. In this case, the total Rubisco flux is RBC (1 + 10^φ^). After computing each phase plane, its y-axis was rescaled to show the total Rubisco flux (instead of just the carboxylation alone). For more details regarding the modelling of Rubisco activity, see the Jupyter notebook “rubisco_phase_planes.ipynb” in our GitHub repository.

### Strains and plasmids used in this study

All strains, plasmids, and gene variants used in this study are listed in **Supplementary Data 1**. Two base strains, both derivative of *E. coli* SIJ488 (Addgene #68246), were employed for growth-coupling experiments. The Δ*rpi* strain was used to enforce the detoxification activity of Rubisco (47). In contrast, the GLY-AUXΔ*gcvTHP* (GLY-AUX2) strain, derivative of the previously described GLY-AUX (38), was created to rely on the product of the oxygenase reaction of Rubisco (2-phisphoglycolate) to support glycolate biosynthesis and complementing the auxotrophy. The *gcvTHP* operon was deleted using the integrated lambda-red system in the SIJ488 strain(66).

### Construction of the pCBB2 plasmid

The pCBB plasmid (p15A ori and *cm^R^*) and the Rubisco sequences used in this study were kindly donated by Ron Milo’s lab (45, 51, 67). The pCBB2 plasmids were designed to allow replacement of Rubisco genes or *prk* using Gibson or USER cloning (see **Supplementary Table 1** for primers list). The *prk* used for creating the first variants of pCBB2 comes from the gene harbored in Chromosome 2 of *C. necator* (prk1). The carbonic anhydrase gene present in pCBB was removed. pCBB2 plasmid architecture was obtained by employing a pSEVA221(50) (RK2 ori and *kan^R^*), cloned in its cargo with a Pj5 promoter (48, 49) upstream of rbc4 and prk1. The ribosome binding sites (RBSs) for the two coding DNA sequences (CDSs) in pCBB2 were determined using the RBS calculator from the Salis lab (68), which provided the translation initiation rate (TIR) values reported in this study. All plasmids were transferred to the *E. coli* selection strains Δ*rpi* and GLY-AUX2 through electroporation.

### Routine strain cultivation

For routine strain handling, lysogeny broth (LB) medium (composed of 1% NaCl, 0.5% yeast extract, 1% tryptone) was used. When appropriate, antibiotics (kanamycin (50 μg/mL), ampicillin (100 μg/mL), or chloramphenicol (25 μg/mL)) were added. Incubation occurred at 37 °C. Then, single colonies were inoculated into a 14 mL clear round-bottom Grenier tubes containing 4 mL of liquid LB medium, supplemented with the required antibiotics, and incubated at 37 °C in a shaking incubator (250 rpm). Once the cultures reached a cell turbidity above 1.0 OD_600_, they were diluted 1% (v/v) into M9 minimal medium supplemented with the required carbon sources for “relaxing” or “selective” conditions (see below).

For growth tests, we used M9 minimal medium (50 mM Na_2_HPO_4_, 20 mM KH_2_PO_4_, 20 mM NH_4_Cl, 2 mM MgSO_4_, 1 mM NaCl, 134 μM EDTA, 100 μM CaCl_2_, 13 μM FeCl_3_·6H_2_O, 6.2 μM ZnCl_2_, 1.62 μM H_3_BO_3_, 0.76 μM CuCl_2_·2H_2_O, 0.42 μM CoCl_2_·2H_2_O, 0.081 μM MnCl_2_·4H_2_O). Then, carbon sources were added to the medium as follows: for GLY-AUX Δgcv, the “selective” medium (M9S) included 20mM glycerol and 5mM formate, while the “relaxing” medium (M9R) additionally includes 2mM glycolate; for Δ*rpi*, the M9S medium included 20 mM glycerol. No M9R exists for this strain, since Δ*rpi* does not confer an auxotrophic phenotype. If preculturing occurs with strain harboring plasmids, antibiotics are added accordingly: chloramphenicol (25 μg/mL) for pCBB, and kanamycin (50 μg/mL) for pCBB2. When growing Δ*rpi* strain harboring any of the pCBB plasmids, plates and tubes were stored in incubators with enriched CO_2_ atmosphere (pCO_2_ = 0.1 atm, corresponding 10% (v/v) CO_2_).

### *In vivo* growth-coupling experiments using Δ*rpi* and GLY-AUX2

Once the precultures on minimal medium have reached an OD_600_ ∼ 0.5 the plate experiment could start. Only for strains harboring any of the pCBB variants, preculturing occurred in M9S and dense cultures were transferred a second time to ensure that there was no carryover from LB or M9R media.

For the plate reader experiment, once the precultures have reached an OD_600_ of ∼ 0.5, 1mL of the grown cultures was harvested by centrifugation at 10,000 × g for 30 s and washed three times with M9 minimal medium without any carbon source. Then, the washed cells were diluted to a final OD_600_ of 0.01 in a final volume of 2 mL (M9S or M9R) and aliquoted in 96-well microtiter plates (Nunclon Delta Surface, Thermo Scientific), such that each well contained 150 μL of cultivation media and 50 μL mineral oil (Sigma-Aldrich) to prevent evaporation while allowing gas exchange. A BioTek Epoch 2 plate reader (BioTek, Bad Friedrichshall, Germany) was used to monitor the growth of technical duplicates at 37 °C by measuring the absorbance (at 600 nm) every 10 min with intermittent orbital and linear shaking, using the same program as previously described (38). When a CO_2_ enriched environment was required, a BioTek Synergy H1 (BioTek, Bad Friedrichshall, Germany) equipped with a programmable CO_2_ control unit was used. Finally, inferred growth rates and final OD_600_ values were used to estimate the capacity of the Prk-Rubisco module variants. These post-run analyses were performed using QurvE (69) and the growth2fig script (36).

When performing growth complementation experiments in baffled shake-flasks for GLY-AUX2, precultures grown on M9R with the corresponding antibiotic were transferred into 50 mL of M9S liquid medium in 250 mL Duran^®^ baffled flask (Merck) sealed with sterile screw caps. Cells were washed three times using M9S media to minimize the effect of carryover and their initial OD_600_ was set at 0.01. We considered a value ≤ 0.05 (corresponding to at least two doublings) as no growth. The flasks, sealed to prevent evaporation of the medium, were placed in shaking incubators at 37 °C and 250 rpm, and cell turbidity was monitored at regular intervals of 72 hours until cultures reached the stationary phase.

### Adaptive laboratory evolution of GLY-AUX2 and pCBB2 in a turbidostat

ALE campaigns were performed in Pioreactors, consisting of 60 mL borosilicate glass vials (Macherey-Nagel) with a working volume of 20 mL, operated in turbidostat mode using the Pioreactor control software. Cultures were maintained at 37 °C with stirring at 750 rpm, and OD_600_ measurements were recorded every 5 min. Turbidostat dilutions were triggered once cell turbidity reached 60% of the maximum OD_600_ observed in batch mode, exchanging 20–90% of the working volume (4–18 mL) per event to maintain mid-exponential growth throughout the campaign. The time interval between dilution events was used as a real-time proxy for population growth rate improvement.

Four reactors (R1–R4) were operated under oxygenation-selective conditions (M9 minimal medium supplemented with 20 mM glycerol and 5 mM formate) for five weeks (∼850 h). Two additional reactors (R5–R6) were run in parallel under relaxed conditions (M9 supplemented with 20 mM glycerol, 5 mM formate, and 2 mM glycolate) with GLY-AUX2 harboring empty pSEVA221, to enable identification and filtering of mutations arising from genetic drift or adaptation to the cultivation platform independently of oxygenation-selective pressure.

Upon emergence of a stable evolved phenotype, chromosomal DNA was extracted from each population for whole-genome sequencing. Plasmid DNA was isolated independently from R1–R4 and sequenced. Identified mutations were retroengineered into naïve pCBB2rbc4prk1 for functional validation.

### Sequencing of the evolved strains

Post-ALE sequencing was performed on reactor populations, isolated clones, and the parental strain. Samples were subjected to whole-genome sequencing (WGS) using a hybrid approach combining Oxford Nanopore long-read sequencing and Illumina short-read sequencing to generate high-quality genome assembly scaffolds. Data analysis was performed using Geneious Prime (version 2025.2.2).

For population sequencing, variants were identified using the “Find SNPs/Variations” function with a minimum variant frequency threshold of 70%. Only mutations present in ≥70% of the population were considered propagating mutations. This approach was used to assess mutations occurring on the chromosome when comparing populations evolved under selective versus relaxed conditions.

To exclude pre-existing variants, all mutations detected in the parental strain were identified and removed from subsequent analyses. This accounted for genomic deletions and sequence differences relative to the *E. coli* SIJ488 reference genome (66). Furthermore, mutations detected in both selectively evolved and relaxed-condition turbidostat populations were excluded, as these were considered unrelated to selective pressure for growth via the Rubisco oxygenation reaction.

To assess mutations occurring on the pCBB2 plasmid, plasmid DNA was isolated from populations R1–R4 and subjected to whole-plasmid sequencing using the same hybrid approach. As control populations grown under relaxed conditions (R5 and R6) carried an empty pSEVA221 plasmid instead of pCBB2, no direct plasmid-level comparison was possible. Therefore, all mutations detected in pCBB2 were reported without background filtering against relaxed-condition controls.

Finally, isolated single clones from populations R1–R4 were sequenced prior to proteomic analysis. All detected SNPs were recorded. As these represent single isolates rather than mixed populations, their mutation patterns may differ from those observed in the corresponding population-level sequencing data.

### Proteomics

All GLY-AUX2 strains were cultivated in 4.5 mL of M9R at 37 °C shaking at 250 rpm in quadruplicates. At OD_600_= 0.5 (mid-exponential phase), cells were harvested at an equivalent of 1 mL OD_600_ = 1.0 and washed with ice-cold PBS (12 mM phosphate buffer, 2.7 mM KCl, 137 mM NaCl, pH = 7.4) before flash-freezing the pellets in liquid nitrogen for storage at −70 °C.

To isolate the proteome, pellets were resuspended in 200 µL freshly prepared ice-cold lysis buffer (6 M urea, 10 mM ammonium bicarbonate, 2 mM MgCl_2_, 20 mM Tris-HCl pH 7.0, 125 U/mL Benzonase in water) and lysed by bead-beating with 150 mg 0.1 mm zirconium beads in a Spex GenoGrinder (3 × 5 min at 1500 rpm with 5 min on ice between cycles). Samples were reduced and alkylated using DTT (20 µL, 55 mM) and iodoacetamide (20 µL, 120 mM), diluted with 450 µL 100 mM ammonium bicarbonate, and 500 µL per sample were digested using 2 µg Trypsin/LysC (Promega, Cat#V5072) at 37°C for 17 h. Digestion mixture was cleaned-up using MCX 2mg Sorbens plates (Oasis MCX 96-Well) by loading 500 µL digest, washing with 2% formic acid and 5% methanol, and eluting with 2% ammonium hydroxide in 40% acetonitrile, followed by dilution with 0.1% formic acid to a final volume of 100 µL. An equivoluminal pool of all samples was prepared as technical control during MS measurements. Peptide concentration was measured using a fluorimetric peptide assay kit (Thermo Scientific, Cat#23290).

Peptide mixtures were analyzed by LC-MS on a Bruker timsTOF HT mass spectrometer coupled to an Agilent 1290 Infinity II LC system operated at analytical flow. Peptides (5 µg per injection) were separated on a Waters CSH C18 column (1.7 μm, 100 Å, 30 × 2.1 mm) heated to 50 °C at a flow rate of 0.5 mL/min using 0.01% formic acid in water (solvent A) and 0.01% formic acid in acetonitrile (solvent B). The separating gradient was 3% to 36% solvent B over 5 min, followed by an increase to 80% B at 0.8 mL/min for 0.5 min, a 0.2 min high-organic wash, and re-equilibration to starting conditions for 2 min. DIA-MS data were acquired in diaPASEF mode using a VIP-HESI electrospray source (Bruker VIP-HESI, Bruker Daltonics; 3.0 kV capillary voltage, nebulizer 4.5 bar, drying gas 10.0 L/min at 240 °C, sheath gas 4.8 L/min at 450 °C). The diaPASEF method sampled an ion mobility range from 1/K0 = 1.30 to 0.70 Vs/cm² with 72 ms ion accumulation and ramp time in the dual TIMS device, over an m/z range of 400-1125 and a total cycle time of ∼0.7 s. Collision energy was ramped from 59 eV at 1/K00 = 1.6 Vs/cm² to 20 eV at 1/K00 = 0.6 Vs/cm², and the TIMS device was linearly calibrated using Agilent ESI-L tuning mix ions (m/z 622.0290, 922.0098, and 1221.9906 with their corresponding 1/K00 values).

Proteins were identified using DIA-NN version 2.2.0 with the default settings using an in-silico spectral library generated from the *E. coli* UniProt reference proteome UP000000625 (downloaded 16. Oct 2025). Tryptic digestion was defined with one missed cleavage, N-terminal methionine excision and fixed carbamidomethylated cysteines, with match-between-runs enabled and precursor-level FDR controlled at 1%. From each sample, outlier replicate was removed and analysis continued with three replicates. Differential expression was identified using FragPipe analyst (70) with settings: protein intensities required 0% non-missing values, significance defined as adjusted p-value < 0.05 and absolute log₂ fold change > 1. Data were median-centered and normalized using a Perseus-type workflow, following correction for multiple testing using the Benjamini-Hochberg method. KEGG pathway enrichment analysis was performed in RStudio (R version 4.5) using the clusterProfiler package (v4.16.0). Gene annotations were obtained from org.EcK12.eg.db (v3.21.0) and pathway data from KEGG REST API using KEGGREST (v1.48.1). Volcano plots were generated using ggplot2 (v3.5.2).

### Retroengineering of ALE mutation and screening in Δ*rpi* and GLY-AUX2

To reintroduce mutations identified in the Prk-Rubisco module during the ALE experiment (M115I in rbc4 and N216T in prk1), USER cloning was performed using the corresponding naïve pCBB2 plasmid as template. USER primers (**Supplementary Table 1**) were designed to introduce the exact single-nucleotide substitutions observed in the ALE strains.

The single mutations rbc4* (M115I) and prk1* (N216T) were introduced via a two-fragment USER assembly. In addition, a three-fragment USER assembly was performed to generate a plasmid harboring both mutations. This resulted in three pCBB2 variants: one carrying M115I, one carrying N216T, and one carrying both mutations.

All assembled plasmids were transformed into chemically competent *E. coli* DH5α by heat shock, and verified clones were stored as cryostocks at −70 °C. Plasmids were subsequently purified and electroporated into the GLY-AUX2 selection strain. Strains were revived by serial culturing in selective medium supplemented with kanamycin, and growth was assessed in microtiter plate reader experiments. Maximum specific growth rate and maximum OD_600_ were determined for each variant and compared to the parental pCBB2 plasmid to evaluate the individual and combined effects of the mutations on flux through the CBB module.

### *In vitro* analysis of Rubisco’s *k_cat,C_*

The catalytic rate of the carboxylation of the mutant Rubisco was measured *in vitro* using a coupled assay from cellular extracts as previously described (59, 71). For Rubisco variants expression, BL21(DE3) cells were transformed with either the pCBB2rbc4prk1 or pCBB2rbc4*prk1 plasmid. Transformed cells were grown in 8 ml LB medium supplemented with 50 μg/mL kanamycin in 24-deep-well plates at 37°C, 250 rpm. When cultures reached an OD_600_ of approximately 0.8, incubation was continued at 25°C for 24-48 h. Cells were then harvested by centrifugation (15 min; 4,000 g) and pellets were lysed in 70 µl BugBuster® ready mix (Millipore) for 25 min at room temperature. Lysates were then centrifuged for 30 min at 4,000 g, 4°C, and the resulting supernatants containing the soluble protein fraction were collected and stored at 4°C until assays.

Assays were performed as described in previous studies (59, 71). Prior to measurements, soluble fractions were incubated for 15 min with 4% CO_2_ and 0.4% O_2_ for Rubisco activation. The assay relies on the coupling of 3-phosphoglycerate production by Rubisco to NADH oxidation, allowing Rubisco specific activity to be quantified spectrophotometrically in each sample. To minimize oxygenation reaction, the assays were performed in a solution equilibrated at 4% CO_2_ and 0.2% O_2_ in a gas-controlled plate reader (Infinite® 200 PRO; TECAN) at 30°C. Rubisco’s active site concentration was measured in each sample by repeating the assay with increasing concentrations of 2-C-carboxyarabinitol 1,5-bisphosphate (CABP), a competitive irreversible rubisco inhibitor. Because assays were performed directly on soluble protein fractions, Rubisco concentration could not be estimated a priori. An initial assay was therefore performed with the undiluted soluble fraction and, when needed (i. e. when the rubisco was too concentrated to measure a decay of the activity with CABP) the assay was repeated with a diluted soluble fraction.

Carboxylation rates were eventually calculated by dividing the specific activity of each sample by its measured active site concentration. Since the coupled assay used in this pipeline tends to underestimate the rates, rate values were further corrected by applying a multiplicative factor of 2.1. This factor was derived from a log–log comparison between rates obtained using this assay and literature values for 11 Rubisco variants, as described previously (59).

### Sequence and structural analyses of the different Rubisco and phosphoribulokinase variants

Amino acid sequences of Rubisco and Prk homologs were aligned using the Clustal Omega webserver (72), generating percentage identity matrices and phylogenetic distance outputs. The corresponding phylogenetic trees were visualized using the Interactive Tree of Life (iTOL) online server, version 6 (73).

To gain mechanistic insight into the mutations identified by ALE in CbbM from *R. rubrum* and CfxP from *C. necator*, we built atomic models of their predicted three-dimensional structures in the expected oligomeric states (homodimeric and homooctameric, respectively) (54, 74) using the ColabFold suite (version 1.5.2) (75) via the COSMIC2 online gateway. The structures of the wild-type proteins and ALE mutants were superimposed and analyzed using the visualization tool UCSF ChimeraX (version 1.7.1) (76) to determine mutation locations and their specific effects on intramolecular or intersubunit interactions. For Rubisco, substrate transplantation was performed using AlphaFill (77), enabling structural enrichment with either CO_2_ or O_2_ at the active site based on the experimentally determined holostructures of the homologous enzyme from *Galdieria sulphuraria* (PDB entries: 4F0K for CO_2,_ 4F0H for O_2_) (78). Ribulose-1,5-bisphosphate (RuBP) was instead transplanted from the crystallographic structure capturing CbbM in complex with the five-carbon sugar substrate (79). For Prk, structural superimposition with the homologous enzyme from *Synechococcus elongatus* PCC 7942 (PDB entry: 6KEV) enabled the identification of the cognate Ru5P and ATP substrate-binding sites.

## Acknowledgments

The authors thank Ari Satanowski, Henrik Petri, and Natalia Grabarczyk for technical assistance in working with GLY-AUX and GLY-AUX2 strains, and Alessia Boga for technical assistance in retro engineering the pCBB2 plasmid. This work was supported by The Novo Nordisk Foundation grants NNF20CC0035580 and NNF24SA0100980. E.O. acknowledges support from the European Union through the Marie Skłodowska-Curie grant agreement 101065339, as well as from the projects UNMUTE (NNF25OC0100601) and GLYCO2 (NNF25OC0103238). The financial support from The Novo Nordisk Foundation through grants NNF10CC1016517, NNF18CC0033664, and NNF23OC0083631 to P.I.N. is likewise gratefully acknowledged.

## Contributions

E.O. and P.I.N. conceptualized the study. E.N. performed the metabolic modelling. E.O., S.C., F.L., M.H.-W., K.K.,and R.V. designed and constructed the plasmids used in this study. E.O., S.C., M.H.-W., R.V., and F.L. constructed the strains used in this study. E.O., S.C., M.H.-W., R.V., K.K., and F.L. characterized the strains in vivo. M.P. performed the phylogenetic analyses and sequence identity matrix analyses presented in the study. M.H.-W. performed the adaptive laboratory evolution experiment. M.H.-W. and K.K. performed whole-genome sequencing of the evolved strains and identified the resulting mutations. K.K. collected the proteomics samples, which were analyzed by L.M.H., S.N.L., and M.M. B.d.P. performed the in vitro kinetic measurements for Rubisco and estimated the Φ ratios between carboxylation and oxygenation. M.P. and K.K. performed the in silico structural predictions of Rubisco and Prk. E.O. wrote the manuscript with input from all authors. All authors reviewed the manuscript. E.O. and P.I.N. acquired funding. E.O. and P.I.N. supervised the research.

## Ethics declarations

The authors declare no competing interests.

---

**Extended Data Fig. 1.**
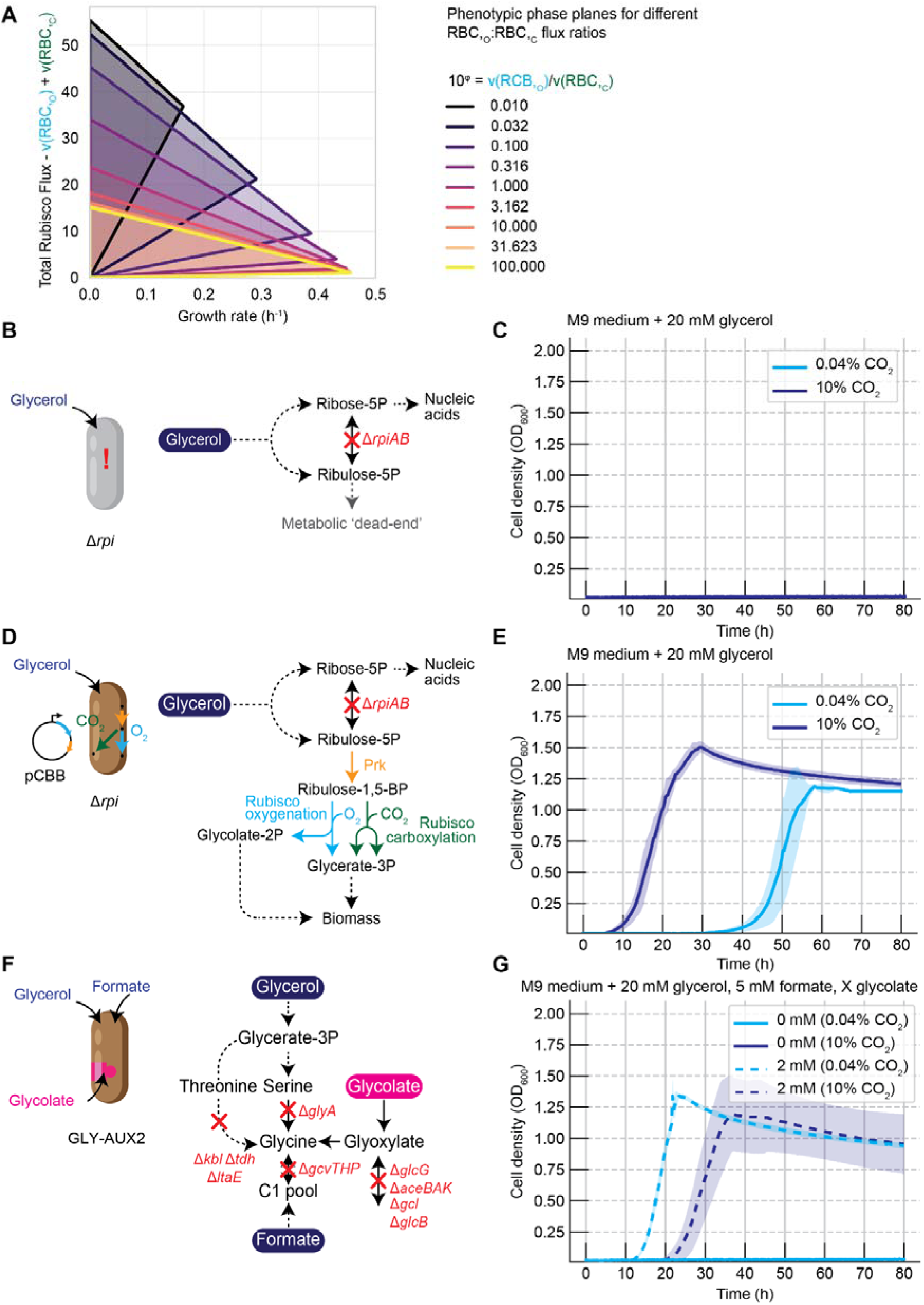
In silico and in vivo validation of the auxotrophic selection designs. A) Phenotypic phase plane analysis of the GLY-AUX2 in silico strain, showing predicted growth rate (x-axis) as a function of total Rubisco flux (carboxylation + oxygenation, y-axis) across different oxygenation-to-carboxylation ratios (10^Φ^). Each shaded area represents a distinct Φ value, illustrating how the model predicts growth complementation across a range of oxygenation contributions. B) Schematic of the Δ*rpi* strain in the absence of pCBB, showing the metabolic dead-end resulting from Ru5P accumulation and the lack of growth complementation without a heterologous Prk–Rubisco module. C) Growth profiles of Δ*rpi* strains without pCBB, showing flat growth curves and confirming the absence of growth complementation under selective conditions. D) Schematic of the Δ*rpi* strain transformed with pCBB, showing growth complementation through Prk–Rubisco activity irrespective of whether carboxylation or oxygenation products are generated. E) Growth profiles of Δ*rpi* strains harboring pCBB under selective conditions at 0.04% and 10% CO_2_. Growth complementation is observed under both conditions, confirming that Δ*rpi* cannot distinguish between carboxylation and oxygenation activities. Curves represent one biological replicate measured in technical triplicate. Shaded areas represent standard error with 95% confidence interval. Experiments were performed in biological triplicate. F) Schematic of GLY-AUX2 showing growth complementation through external glycolate supplementation under relaxed conditions, illustrating the tightness of the selection scheme. G) Growth profiles of GLY-AUX2 under relaxed conditions (2 mM glycolate supplementation) and selective conditions at 0.04% and 10% CO_2_. Growth is restored exclusively under relaxed conditions and at ambient CO_2_, confirming the selectivity of the GLY-AUX2 design for Rubisco oxygenase activity. Curves represent one biological replicate measured in technical triplicate. Shaded areas represent standard error with 95% confidence interval. Experiments were performed in biological triplicate.

**Extended Data Fig. 2.**
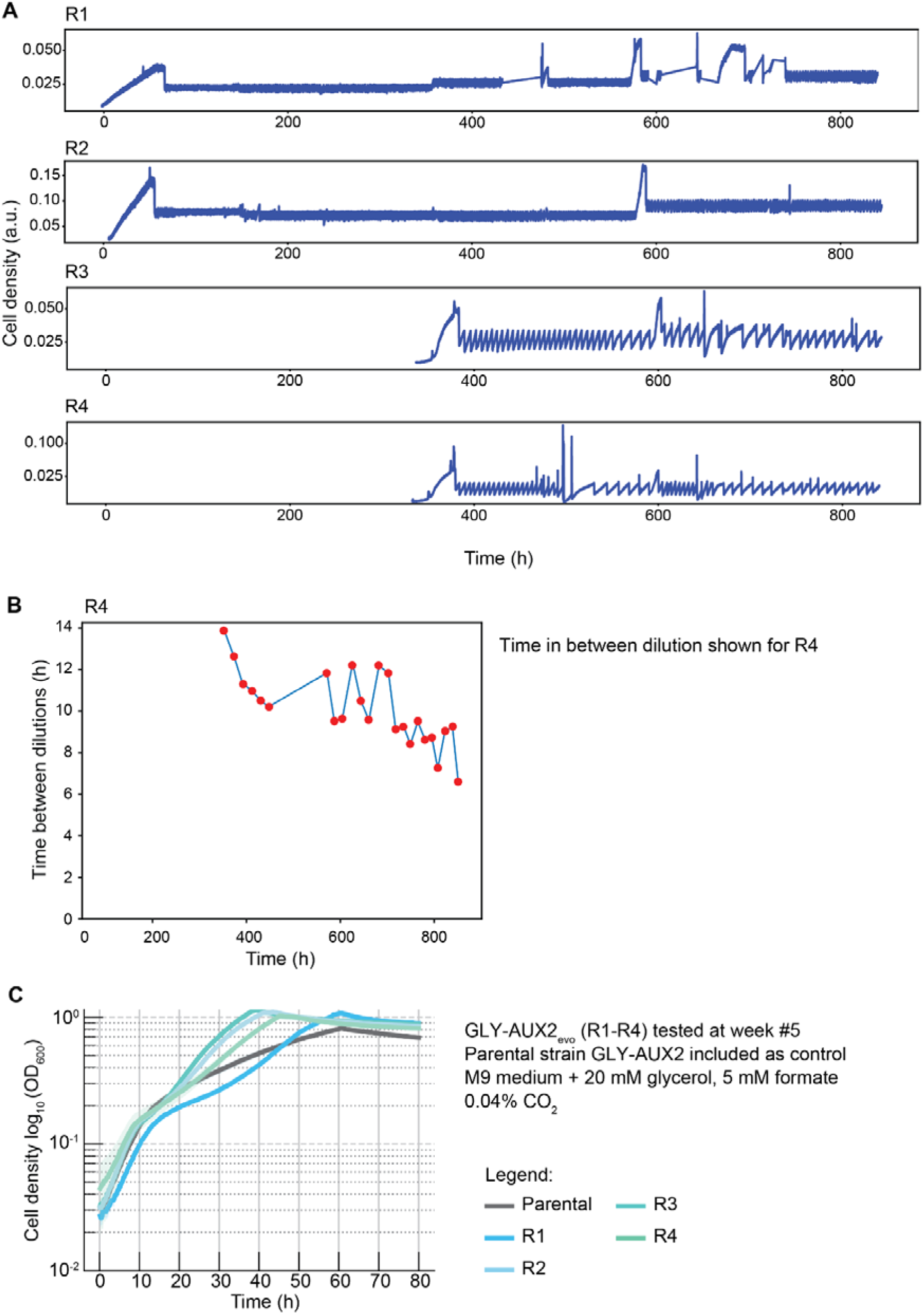
Phenotypic monitoring of the adaptive laboratory evolution campaign. A) Time course of turbidostat runs R1–R4 over five weeks of continuous cultivation, showing cell density (y-axis) as a function of time (x-axis). Each reactor was maintained under oxygenation-selective conditions with automatic dilution to sustain a predetermined turbidity setpoint. B) Schematic illustrating the use of time between dilutions as a proxy for population growth rate improvement during turbidostat cultivation, shown for reactor R4 as a representative example. A decrease in time between dilutions indicates an increase in the maximum growth rate of the evolving population. C) Growth rate comparison between the evolved ALE populations from R1–R4 and the naïve parental GLY-AUX2 strain, measured at ambient CO_2_ under oxygenation-selective conditions. Each dot represents a technical replicate. Experiments were performed on all four evolved populations in technical triplicate.

