## Supplementary for "A microbial growth-coupled platform for *in vivo* interrogation of Rubisco oxygenase activity"

**Authors**

Enrico Orsi^1,*^, Kai Kabuth^1^, Simone Cusimano^1^, Mathias Herløv-Wagner^1^, Rutger Verbakel^1^, Francesco Luppino^1^, Michele Partipilo^1^, Benoit de Pins^2^, Elad Noor^3^, Lena Maria Hümmler^4^, Michael Mülleder^4^, Steffen N. Lindner^4^, Markus Ralser^4^, Pablo I. Nikel^1,*^

**Affiliations**

^1^ BRIGHT, Technical University of Denmark, 2800 Kgs Lyngby, Denmark

^2^ Department of Biology, University of Naples “Federico II”, Naples, Italy

^3^ Department of Plant and Environmental Sciences, Weizmann Institute of Science, Rehovot, Israel

^4^ Department of Biochemistry, Charité Universitätsmedizin Berlin, Freie Universität Berlin and Humboldt-Universität, 10117, Berlin, Germany

*** Correspondence to**

Enrico Orsi

and Pablo I. Nikel

**This PDF file includes:**

Figs. S1 to S12

Tables S1 to S6

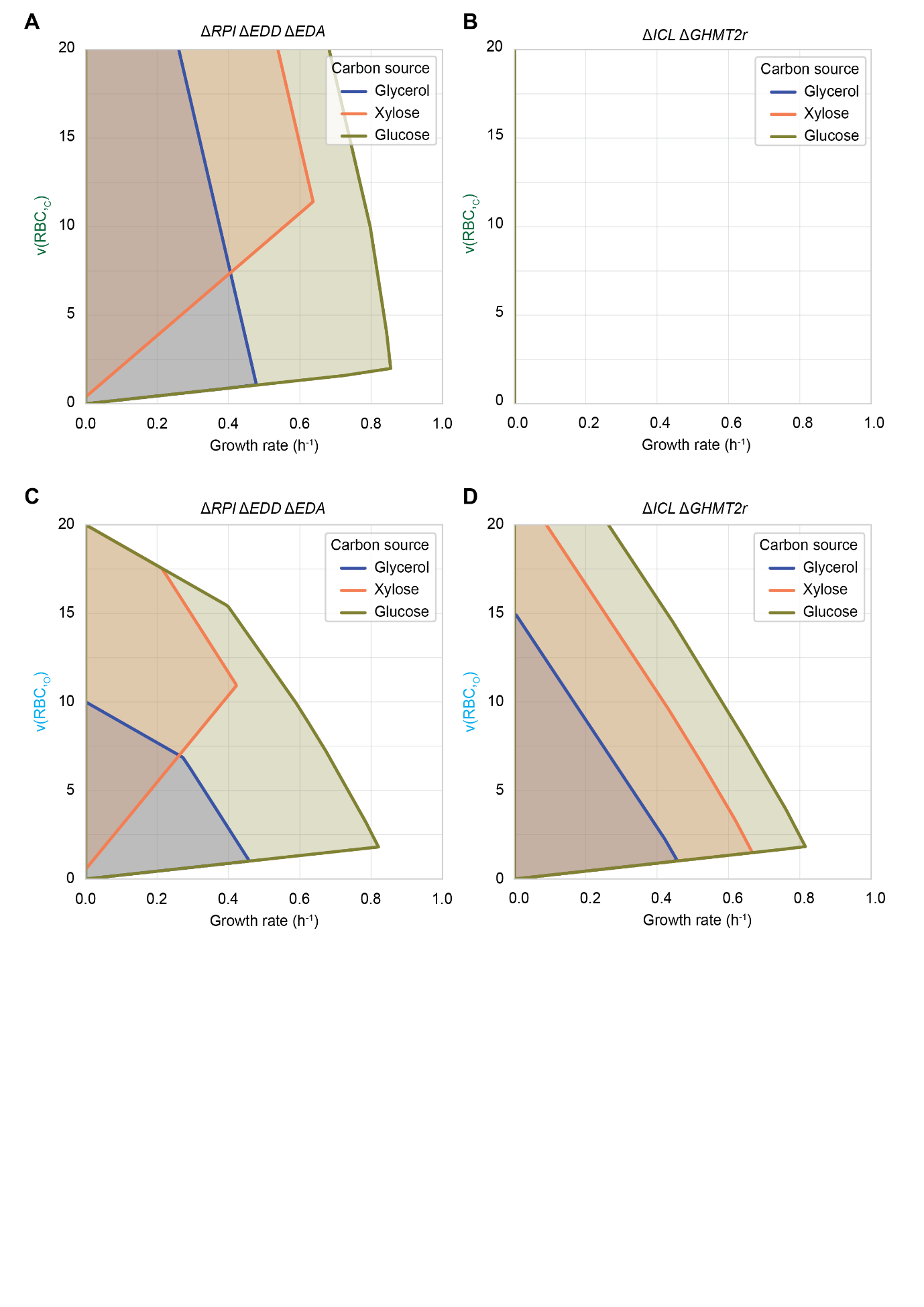

**Fig. S1.**

Phase plane analysis of growth rate as a function of Rubisco carboxylation and oxygenation flux in two selection strains. Predicted growth rate as a function of flux through the carboxylation (v(RBC_,C_); panels A, B) and oxygenation (v(RBC_,O_); panels C, D) reactions in ∆rpi (panels A, C) and GLY-AUX2 (panels B, D). The modelling was performed on a compact *E. coli* metabolic model (1).

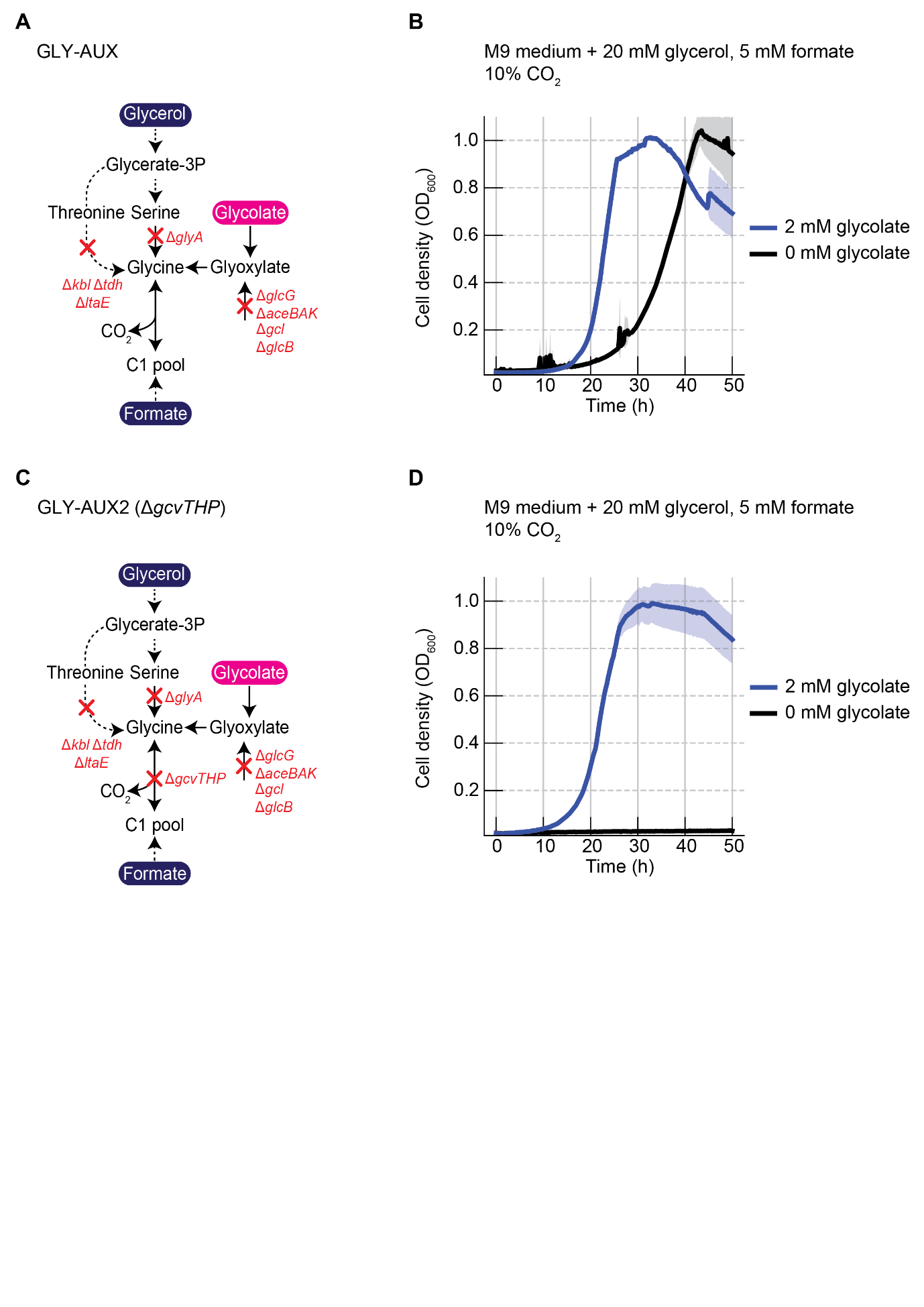

**Fig. S2.**

Overview of GLY-AUX and GLY-AUX2 when growing at 10% CO_2_ (pCO₂ = 0.1 atm). Deletion of the glycine cleavage system complex (*gcvTHP*) prevents CO_2_ and formate to be converted into glycine. The growth profiles represent the average of one biological replicate measured as technical triplicate.

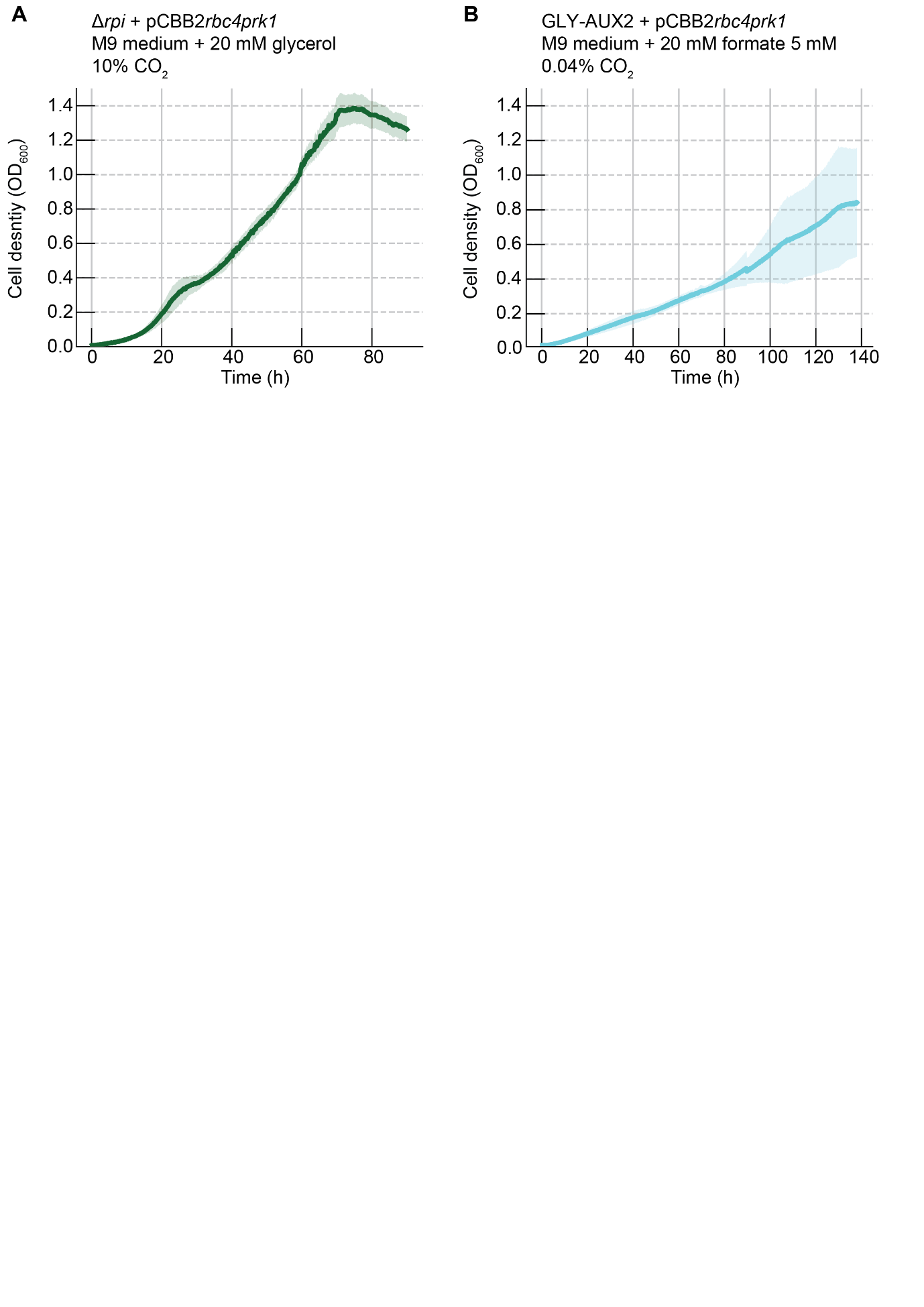

**Fig. S3.**

Growth profiles of the ∆*rpi* and GLY-AUX2 strains transformed with pCBB2 and cultivated in M9 medium under selective conditions. The pCBB2 plasmid harbors the *cbbM* gene from *Rhodospirillum rubrum* (*rbc4*), and the *prk* gene from the chromosome 2 of *Cupriavidus necator* (*prk1*). Characterization of the ∆*rpi* strain was performed at 10% CO_2_ (pCO_2_ = 0.1 atm) to use growth as a proxy for carboxylation activity, whereas for the GLY-AUX2 strain, growth was monitored at ambient CO_2_ concentrations (0.04% CO_2_, equivalent to pCO_2_ = 0.0004 atm) to assess oxygenase activity. The curve shown is a representative growth profile from three biological replicates; the halo around each line represents the standard error around each data point, with a 95% confidence interval. For details on the architecture of pCBB2, consult the main text and the plasmid maps deposited in Supplementary Data 1.

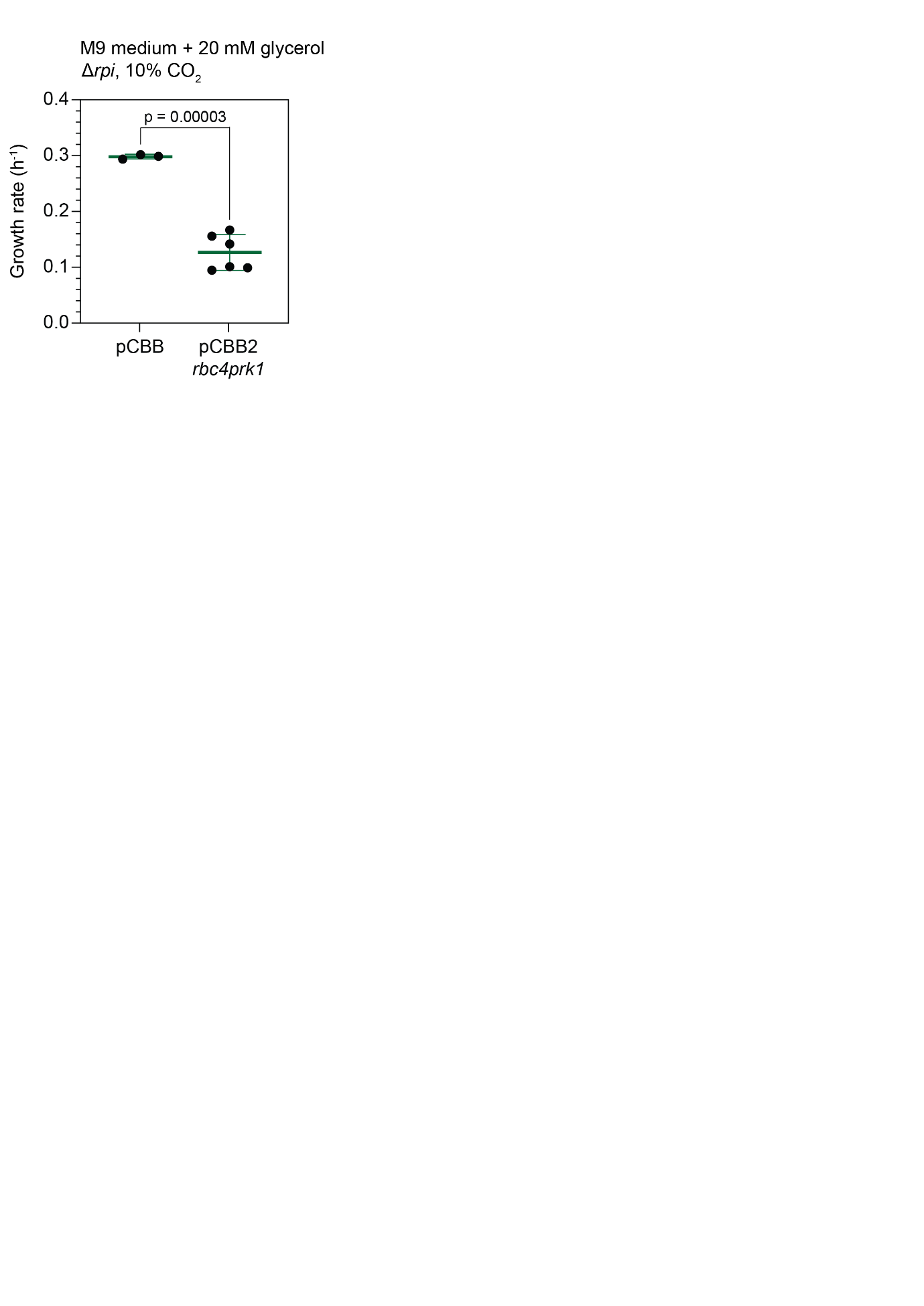

**Fig. S4.**

Comparison of growth rates between pCBB and pCBB2*rbc4prk1* in ∆*rpi* at 10% CO_2_. Growth rates of ∆*rpi* strains harboring pCBB or pCBB2*rbc4prk1*, cultivated at 10% CO_2_. Data are mean ± s.d. (n ≥ 3 biological replicates); individual points represent biological replicates. Statistical comparison by two-tailed unpaired t-test, p-value reported in the figure.

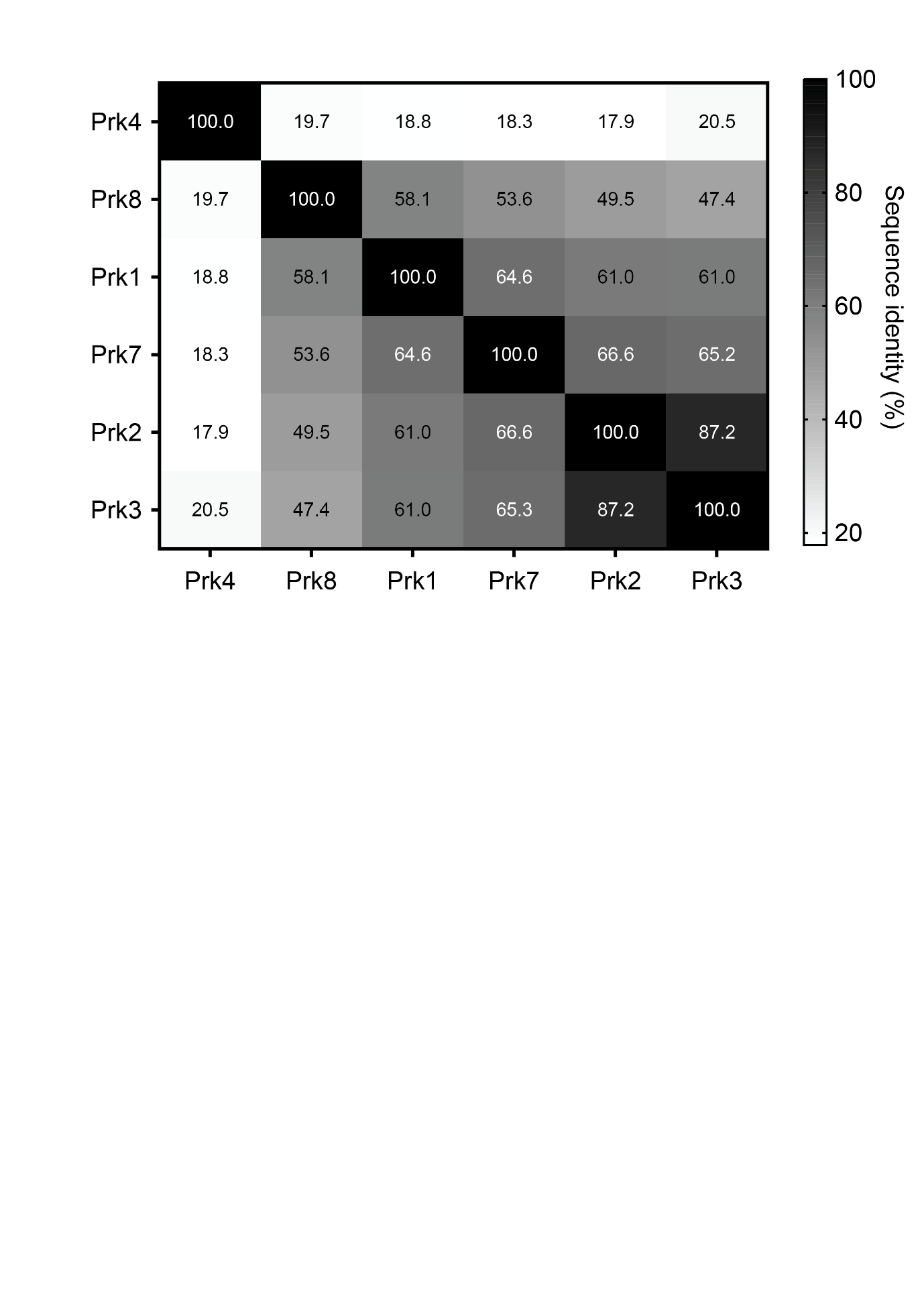

**Fig. S5.**

Percentage identity matrix of phosphoribulokinase variants used in this study. Pairwise sequence identities among the eight Prk variants were calculated from amino acid sequence alignments using Clustal Omega (2). Protein names and source organisms are listed in Supplementary Data 1.

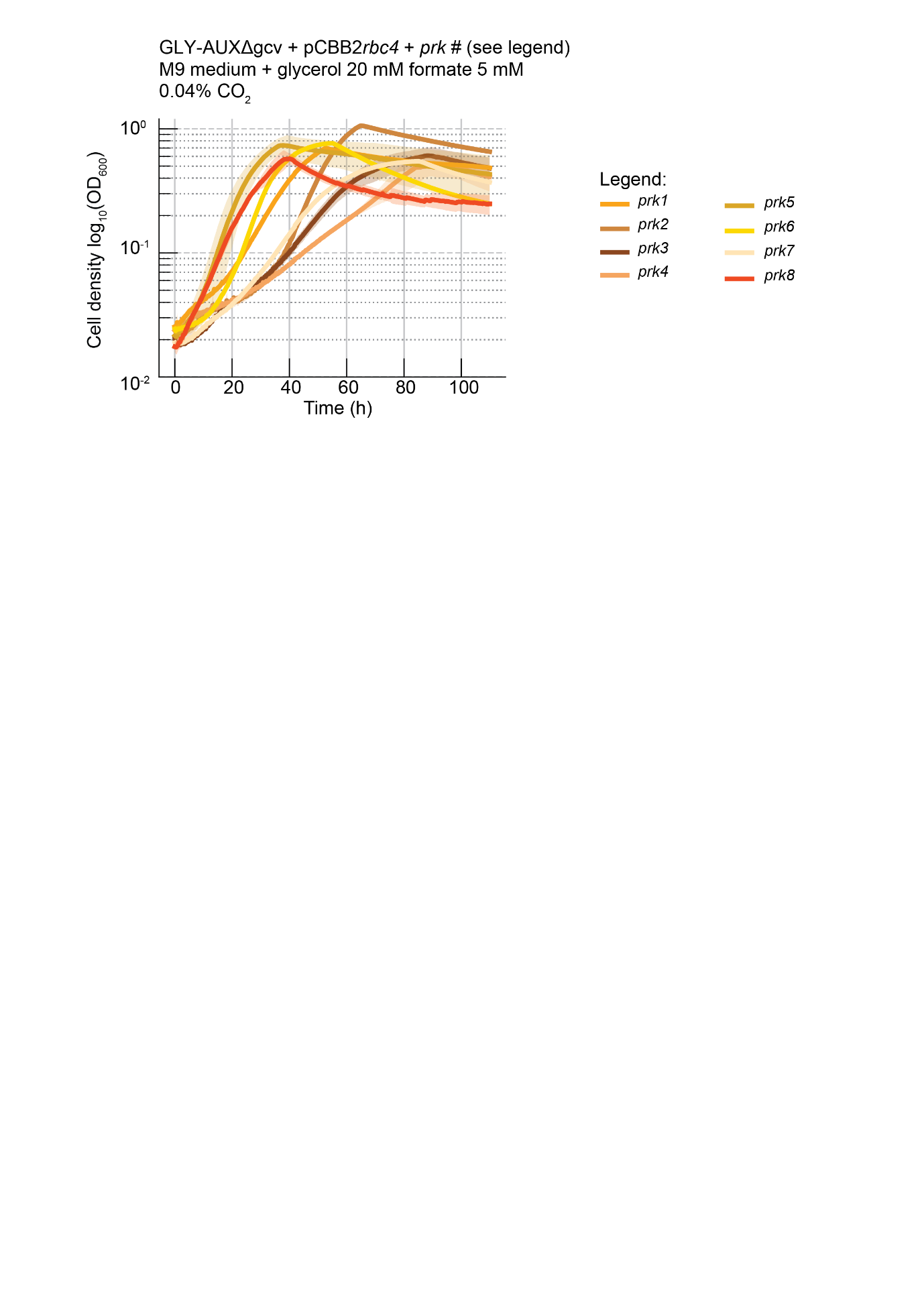

**Fig. S6.**

Growth profiles of GLY-AUX2 strains harboring pCBB2 variants with different Prk variants under oxygenation-selective conditions. Strains were cultivated in M9 medium at ambient CO2 (0.04%, pCO2 = 0.0004 atm). All constructs carry *rbc4* (*cbbM* from *Rhodospirillum rubrum*) paired with the indicated *prk* variant (see legend and Supplementary Data 1). Curves shown are representative growth profiles from one biological replicate per variant, measured in technical triplicate. Shaded areas represent standard error with 95% confidence intervals.

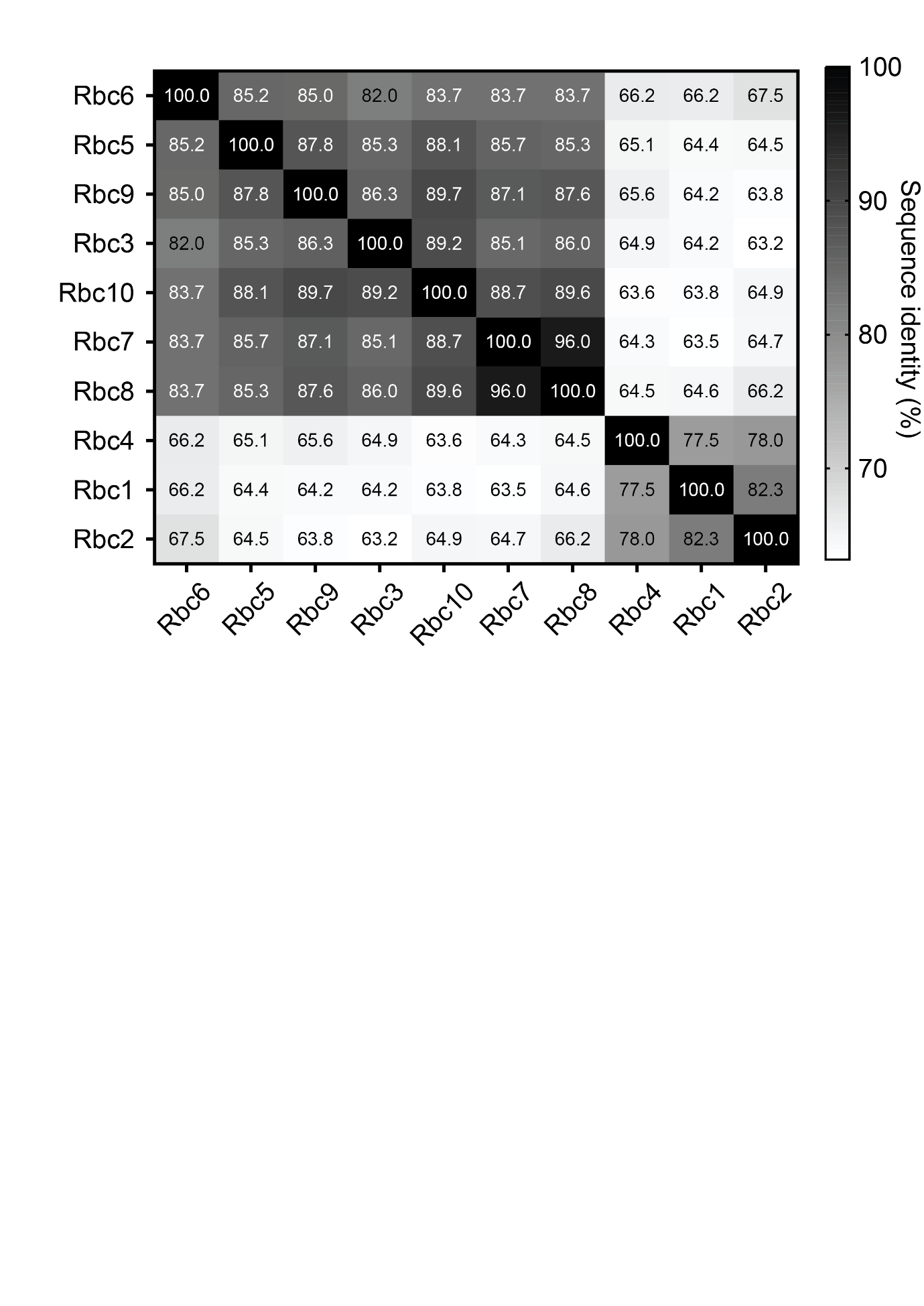

**Fig. S7.**

Percentage identity matrix of Form II Rubisco variants used in this study. Pairwise sequence identities among the ten Rubisco variants were calculated from amino acid sequence alignments using Clustal Omega (2). Protein names and source organisms are listed in Supplementary Data 1.

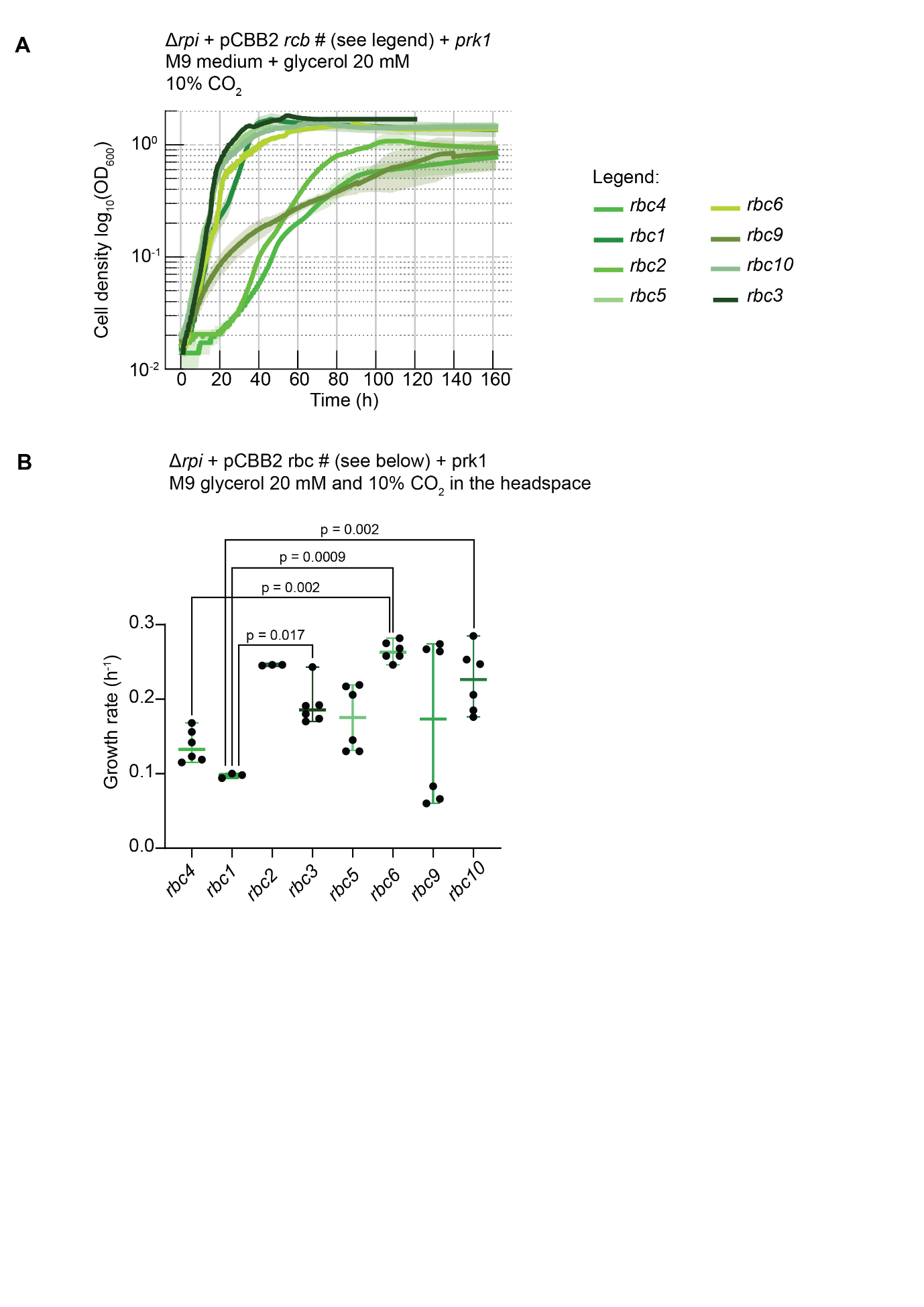

**Fig. S8.**

Growth complementation of different rubisco variants in the ∆*rpi* strain. A) Growth profiles of ∆*rpi* transformed with pCBB2 and cultivated in M9 medium under selective conditions. The pCBB2 plasmid variants harbor *prk1* and different Form II Rubisco variants. Growth was monitored at 10% CO_2_ (equivalent to pCO_2_ = 0.1 atm) to assess the carboxylation capacity of each plasmid. The curves shown are representative growth profiles from one biological replicate per *cbbM* variant, measured in technical triplicate. B) Growth rates of ∆*rpi* strains harboring pCBB2*prk1* variants carrying different *rbc* genes at 10% CO_2_. Individual points represent biologically independent experiments (n = 3–6); bars indicate mean ± s.d. Gene names and source species are listed in Supplementary Data 1. Statistical comparisons by one-way ANOVA with Tukey's multiple comparisons test; significant p-values are indicated in the figure.

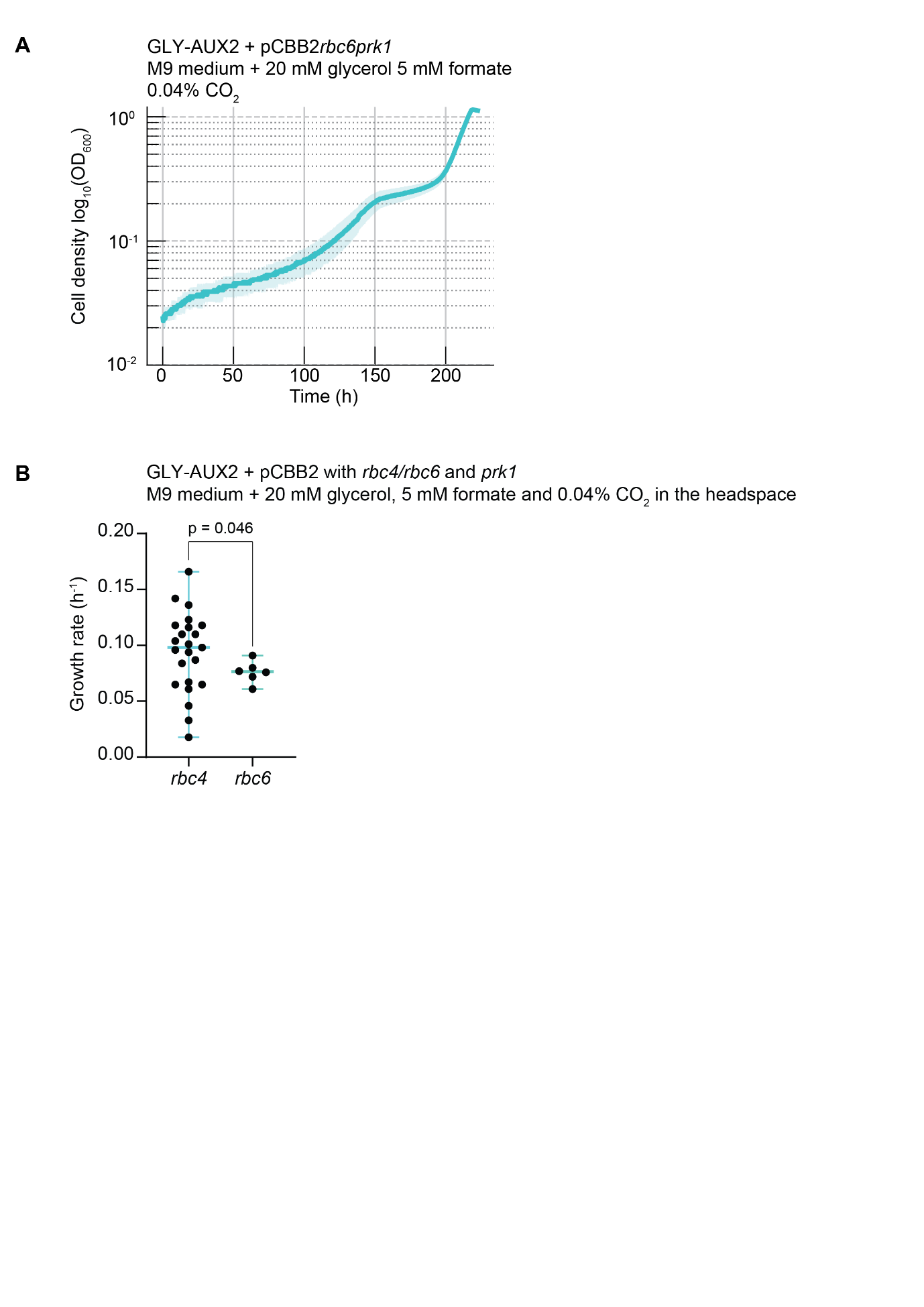

**Fig. S9.**

Growth complementation of GLY-AUX2 with pCBB2 variants in a plate reader. A) Growth profile of GLY-AUX2 transformed with pCBB2 harboring *prk1* and *rbc6* (*cbbM* from *Sulfurivirga caldicuralii*), cultivated in M9 medium at ambient CO_2_ (pCO_2_ = 0.0004 atm) under oxygenation-selective conditions. The curve is demonstrative and represents one biological replicate measured in technical triplicate. Shaded area represents standard error with 95% confidence interval. B) Inferred growth rates of GLY-AUX2 strains harboring pCBB2 variants detectable in the plate reader setup. Data are mean ± s.d. (n ≥ 3 biological replicates); individual points represent biological replicates. Statistical comparisons by two-tailed unpaired t-test. Gene names and source organisms are listed in Supplementary Data 1.

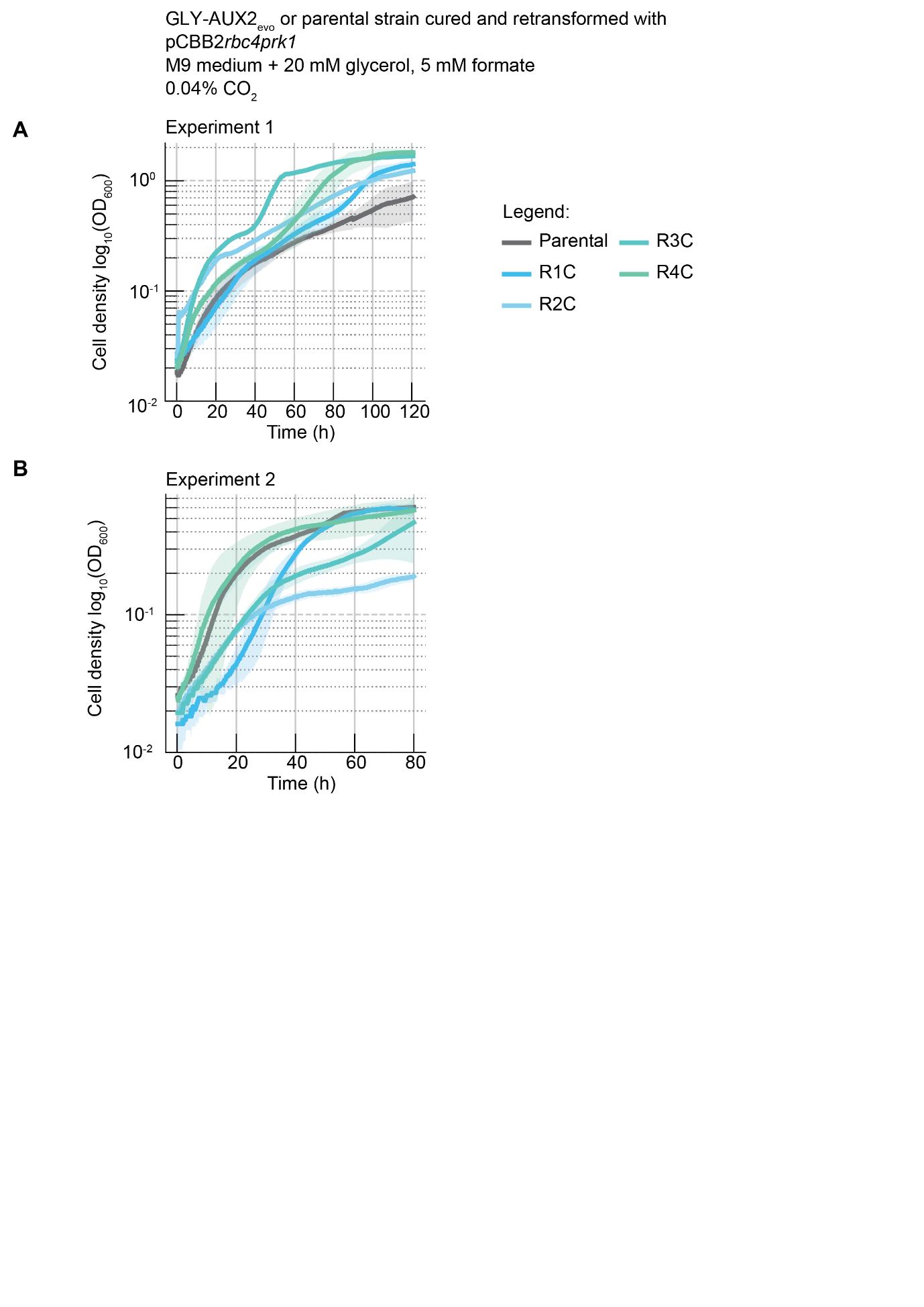

**Fig. S10.**

Growth complementation of GLY-AUX2_evo_ isolates retransformed with naïve pCBB2*rbc4prk1*. Growth of GLY-AUX2_evo_ clones isolated from ALE reactors R1–R4 and retransformed with naïve pCBB2*rbc4prk1*, compared to the parental GLY-AUX2 strain harboring the same plasmid. Growth was monitored at ambient CO_2_ (0.04%, pCO_2_ = 0.0004 atm) under oxygenation-selective conditions. Curves are representative growth profiles from one biological replicate per isolated population, measured in technical triplicate, from two independent experiments.

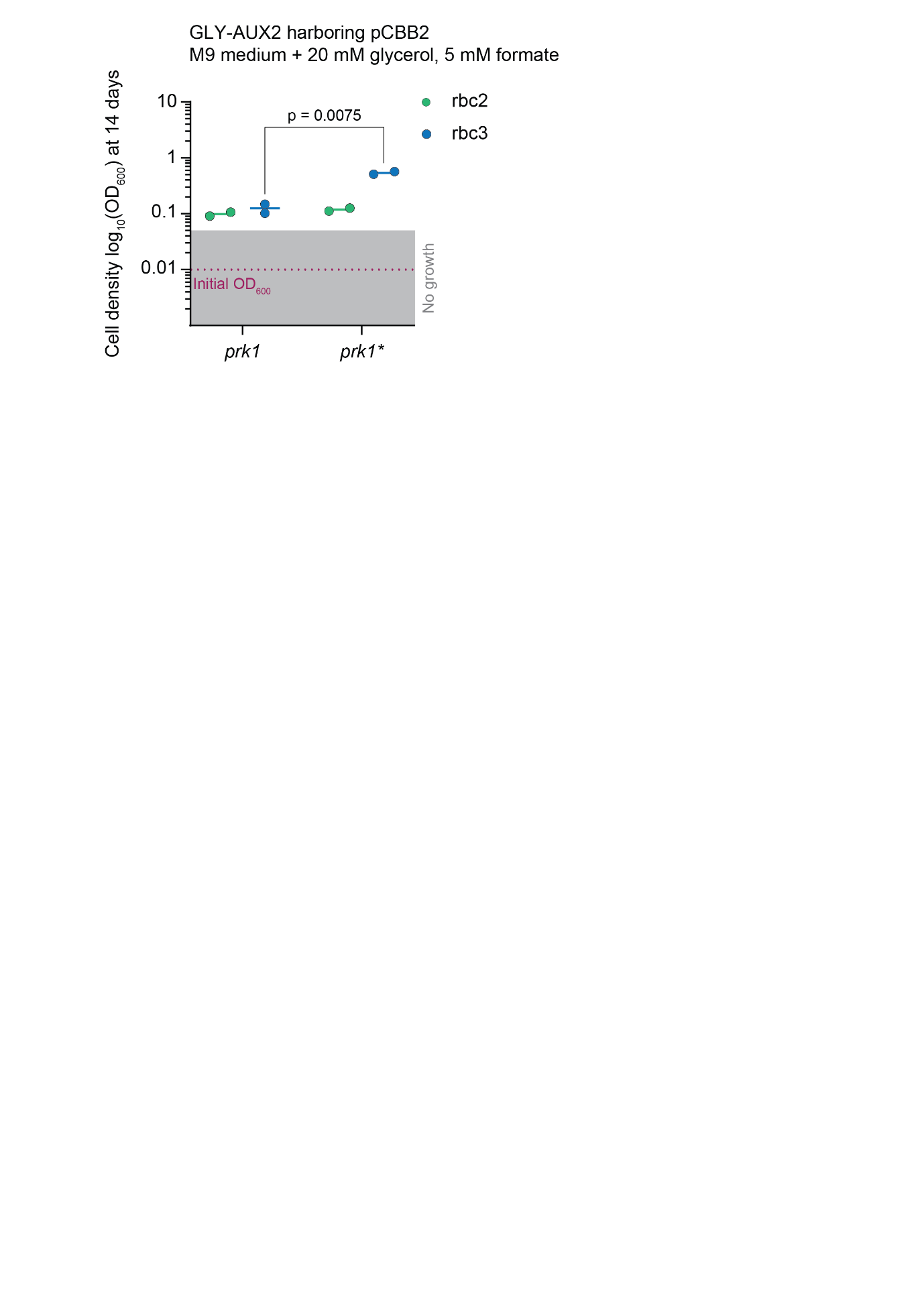

**Fig. S11.**

Effect of the *prk1* mutation on oxygenation-dependent growth of *rbc2* and *rbc3* variants in shake flasks. Final OD_600_ values after 14 days of incubation in sealed baffled shake flasks under oxygenation-selective conditions. Each dot represents a biological replicate. Statistical significance was assessed using an unpaired two-tailed t-test, with the corresponding p-value = 0.0075.

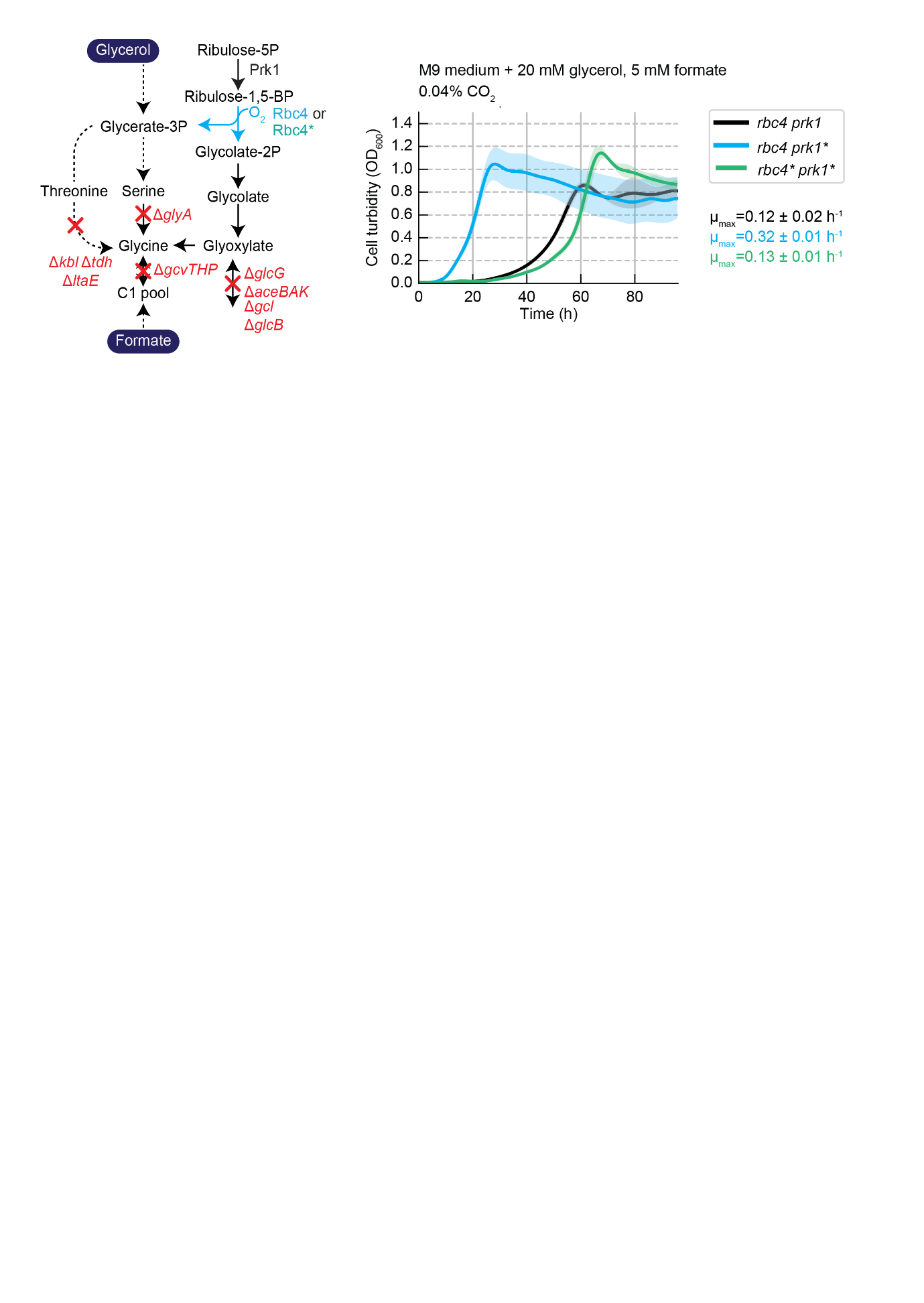

**Fig. S12.**

Effect of *rbc4* and *prk1* mutations on oxygenation-dependent growth of GLY-AUX2 at ambient CO_2_. Growth profiles of GLY-AUX2 strains harboring pCBB2 variants carrying combinations of *rbc4/rbc4** and *prk1/prk1**, cultivated at ambient CO_2_ (0.04%, pCO_2_ = 0.0004 atm). Each line represents the mean of four biological replicates and shaded areas represent standard error with 95% confidence interval. Statistical comparisons were performed using one-way ANOVA followed by Tukey's post-hoc test, revealing that *rbc4prk1** grew significantly differently compared to the other two conditions (adjusted p-value <0.0001), while no significant difference was observed between the remaining two comparisons (adjusted p-value = 0.5981).

**Table S1.**

Primers used in this study. Purpose (1 = cloning of pCBB2, 2 = retro-engineering of mutations in the *cbbM_Rr_* and *cfxP_Cn-C2_* CDSs after ALE)

| **Primer Name** | **Sequence (5' -> 3')** | **Purpose** |
| --- | --- | --- |
| FL06_CBB2.0bb_U_fw | ATGATGGGUCCAGGCATCAAATAAAACGAAAG | 1 |
| FL07_CBB2.0bb_U_rv | ACCGAACUTACGCCGGAAGGGCGCTG | 1 |
| FL18_prk7_U_fw | AGAACGUTATCCCATTATCGCTATCACCGG | 1 |
| FL19_prk7_U_rv | ACGTTCUGACATAGGCGGCTCCGCTTT | 1 |
| FL20_OptRbs_U_rv | AGGCGGCUCCGCTTTAGTTGTGGTGACA | 1 |
| FL21_prk8_U_fw | AGCCGCCUATGTCGGAACGCTATCCGATCA | 1 |
| FL22_prk8_U_rv | ACCCATCAUTACAGGGCTGCGCGCTTA | 1 |
| FL23_prk9_U_fw | AGCCGCCUATGTCAGAACGTTATCCCATTATC | 1 |
| FL24_prk9_U_rv | ACCCATCAUTACTGCGCGGCGCGCTTG | 1 |
| FL30_prkAmut_U_fw | AGCCGCCUATGAGCAAGCCAGATCGTGTTGT | 1 |
| FL31_prkAmut_U_rv | ACCCATCAUTAGACACTAGCGGCGACGGGTGC | 1 |
| FL32_prk3_U_fw | AGCCGCCUATGAGCAAGAAGCATCCCAT | 1 |
| FL33_prk3_U_rv | ACCCATCAUCAGGCCACCTTGCTCTCTC | 1 |
| FL34_prk4_U_fw | AGCCGCCUATGTCGAAGAAACATCCGAT | 1 |
| FL35_prk4_U_rv | ACCCATCAUCACGCGACCTTCGACTCAC | 1 |
| FL36_prk5_U_fw | AGCCGCCUGTGTCGAAGAAATATCCCAT | 1 |
| FL37_prk5_U_rv | ACCCATCAUCAGGCCCGCGCGCG | 1 |
| FL38_prk6_U_fw | AGCCGCCUGTGAGCAAAAAATACCCCATTATTAG | 1 |
| FL39_prk6_U_rv | ACCCATCAUCACGCCCGGGCGCGTCG | 1 |
| FL40_prk10_U_fw | AGCCGCCUATGTCTGAACGCTACCCAATT | 1 |
| FL41_prk10_U_rv | ACCCATCAUTACTGGGCCGCACGCTTGCG | 1 |
| FL42_prk11_U_fw | AGCCGCCUATGTCCATCAAGCACCCCATCAT | 1 |
| FL43_prk11_U_rv | ACCCATCAUCAGCCGGCTCGGCGCTTCCT | 1 |
| FL44_prk12_U_fw | AGCCGCCUATGAGCATCAAGCACCCGATCA | 1 |
| FL45_prk12_U_rv | ACCCATCAUCAGCCGGCGCGTCGTTTGCG | 1 |
| FL46_prk13_U_fw | AGCCGCCUATGTCTGCCAAACATCCGGTCA | 1 |
| FL47_prk13_U_rv | ACCCATCAUTACTCGATTTTCTTTCCTTCCATCAA | 1 |
| FL48_prk14_U_fw | AGCCGCCUATGTCAGCAAAACATCCGGTGATC | 1 |
| FL49_prk14_U_rv | ACCCATCAUTATTCAATCTTTTTACCCTCCATTAAACGC | 1 |
| oEO_526 | agttcggUgtcaccacaactaaagcggag | 1 |
| oEO_527 | atacgaccUccttacttatacctcttgaaat | 1 |
| oEO_528 | aggtcgtaUaatggatcagtcctcgcgttacgcc | 1 |
| oEO_529 | accgaacUtacttgacccctaattcatcccgcc | 1 |
| oEO_530 | aggtcgtaUaatggaccaatcgaaccgttatg | 1 |
| oEO_531 | accgaacUtatgccgcccgctgtaaacga | 1 |
| oEO_532 | aggtcgtaUaatggatcaatcaaaccgctatgc | 1 |
| oEO_533 | accgaacUtatttgtgaacgccgagcttct | 1 |
| oEO_534 | aggtcgtaUaatggaccagagtaatcggta | 1 |
| oEO_535 | accgaacUtatttgtgaacgcccaacttc | 1 |
| oEO_536 | aggtcgtaUaatggaccaatcgaatcgc | 1 |
| oEO_537 | accgaacUtatttgtgaactccaagcttct | 1 |
| oEO_538 | aggtcgtaUaatggaccagtcgaatcgt | 1 |
| oEO_539 | accgaacUtatttatgtactcccagtttctca | 1 |
| oEO_540 | aggtcgtaUaatggaccaatctaatcgg | 1 |
| oEO_541 | accgaacUtatttatgcactcccaactta | 1 |
| oEO_542 | aggtcgtaUaatggaccaatcaaaccgcta | 1 |
| oEO_543 | accgaacUtatttgtgaactccgagtttctc | 1 |
| oEO_544 | aggtcgtaUaatggatcaaagcaaccgctatgc | 1 |
| oEO_545 | accgaacUtatttgtggacgcccagtttct | 1 |
| MW_001 | CTCAATCTTCAATGTTGGCACGC | 2 |
| MW_002 | CGATGCTGGGTTTCGTGGTG | 2 |
| MW_003 | AGGTGGCUGCTGAACCCCCAG | 2 |
| MW_004 | ATACCCUGGTTGTTTCCCATGGTGA | 2 |
| MW_005 | GAGGCTGGCCGTAGGCCGGCCGCGA TGCUGGTGGCT | 2 |
| MW_006 | AGGGTAUAGGCGACGTGGAATACGCC | 2 |
| MW_007 | AGGGGGUGGACGTATCCACGCACGG | 2 |
| MW_008 | ACCCCCUTTATCTCGCGCGAAATCC | 2 |

**Table S2.**

Translation Initiation Rates (TIR) comparisons between Rbc4 and different Phosphoribulokinase gene variants in the context of pCBB2. TIR values were calculated using the RBS calculator software (3).

| Name | TIR Rbc4 | TIR Prk variant | TIR Rbc4/Prk# |
| --- | --- | --- | --- |
| Prk1 | 7431 | 1845 | 4.03 |
| Prk2 | 7431 | 2947 | 2.52 |
| Prk3 | 7431 | 1396 | 5.32 |
| Prk4 | 7431 | 6248 | 1.19 |
| Prk5 | 7431 | 3465 | 2.14 |
| Prk6 | 7431 | 803 | 9.25 |
| Prk7 | 7431 | 1454 | 5.11 |
| Prk8 | 7431 | 109 | 68.17 |

**Table S3.**

Translation Initiation Rates (TIR) comparisons between different Rubisco variants and Prk1 in the context of pCBB2. TIR values were calculated using the RBS calculator software (3).

| Rubisco isoform | TIR Rbc # | TIR Prk1 | TIR Rbc#/ Prk1 | Notes |
| --- | --- | --- | --- | --- |
| Rbc4 | 7431 | 1845 | 4.03 | Construct used for the design of pCBB2 |
| Rbc1 | 12752 | 264 | 48.3 |  |
| Rbc2 | 8896 | 48 | 185.33 |  |
| Rbc3 | 14994 | 166 | 90.33 |  |
| Rbc5 | 11191 | 4339 | 2.58 |  |
| Rbc6 | 41837 | 1071 | 39.06 |  |
| Rbc7 | 43763 | 320 | 136.76 |  |
| Rbc8 | 8777 | 60 | 146.28 |  |
| Rbc9 | 17162 | 80 | 214.53 |  |
| Rbc10 | 18610 | 1023 | 18.19 |  |

**Table S4.**

Frequency of *rbc4* and *prk1* mutations in evolved populations from adaptive laboratory evolution campaigns in reactors R1-R4 after five weeks. Percentages are normalized to the total number of sequencing reads. No additional mutations were identified within the coding sequences (CDS) of either gene.

| Reactor | *rbc4* (M115I) | | Total | *prk1* (N216T) | Total |
| --- | --- | --- | --- | --- | --- |
| R1 | ATG 🡪 ATA | ATG 🡪 ATT |  | AAC 🡪 ACC |  |
|  | 30,30% | 4% | 34,30% | 41,40% | 41,40% |
| R2 |  |  |  | AAC 🡪 ACC |  |
|  |  |  |  | 100%* | 100,00%* |
| R3 | ATG 🡪 ATA |  |  | AAC 🡪 ACC |  |
|  | 100%* |  | 100,00%* | 100%** | 100,00%** |
| R4 | ATG 🡪 ATA |  |  | AAC 🡪 ACC |  |
|  | 89,40% |  | 89,40% | 99,20% | 99,20% |

*data obtained from 8 reads

**data obtained from 5 reads

**Table S5.**

Frequency of *rbc4* and *prk1* mutations in evolved populations from adaptive laboratory evolution campaigns in reactor R2 after nine weeks. Percentages are normalized to the total number of sequencing reads. No additional mutations were identified within the coding sequences (CDS) of either gene.

| Reactor | *rbc4* (M115I) | *prk1* (N216T) |
| --- | --- | --- |
| R2 | ATG 🡪 ATA | AAC 🡪 ACC |
|  | 87,70% | 100% |
